# ABFormer: A Transformer-based Model to Enhance Antibody-Drug Conjugates Activity Prediction through Contextualized Antibody-Antigen Embedding

**DOI:** 10.64898/2026.02.03.703522

**Authors:** Rushi Katabathuni, Viren Loka, Sanjana Gogte, Vani Kondaparthi

## Abstract

Computational screening is increasingly becoming a crucial aspect of Antibody-Drug Conjugate (ADC) research, allowing the elimination of dead ends at earlier stages and concentrating on potential candidates, which can significantly reduce the cost of development. The current state-of-the-art deep learning model, ADCNet, usually considers antibodies, antigens, linkers, and payloads as distinct features. However, this overlooks the complex context of antibody-antigen binding, which is primarily responsible for the targeting and uptake of ADCs. To address this limitation, we present ABFormer, a transformer-based framework tailored for ADC activity prediction and in-silico triage. ABFormer integrates high-resolution antibody–antigen interface information through a pretrained interaction encoder and combines it with chemically enriched linker and payload representations obtained from a fine-tuned molecular encoder. This multi-modal design replaces naive feature concatenation with biologically informed contextual embeddings that more accurately reflect molecular recognition. ABFormer outperforms in leave-pair-out evaluation and achieves 100% accuracy on a separate test set of 22 novel ADCs, while the baselines are severely mis-calibrated. Ablation study confirms that the predictive capability is predominantly driven by interaction-aware antibody-antigen representations, while small-molecule encoders enhance specificity by reducing false positives. In conclusion, ABFormer provides a reliable and efficient platform for early filtering of ADC activity and selection of candidates.

## 1. Introduction

Antibody-drug conjugates (ADCs) are a paradigm-shifting class of therapeutic agents designed to selectively deliver highly potent cytotoxic drugs to malignant cells by chemically conjugating a tumor-specific monoclonal antibody to a small molecule warhead via a specialized linker [1]. Following systemic injection, the antibody moiety of the ADC selectively recognizes and binds to a distinctive antigen expressed on the surface of the tumor cells, contributing to the internalization of the entire complex via receptor-mediated endocytosis into the endosomal pathway. The endosomal trafficking of the conjugate to the lysosome positions the linker within a distinctive intracellular environment that is prone to cleavage by programmed cellular processes, including acidic hydrolysis, enzymatic degradation, or reduction of disulfide bonds [2]. The release of the active cytotoxic warhead into the cytoplasm enables it to bind its molecular target, mainly tubulin microfilaments or genomic DNA, to inhibit critical cellular processes and induce apoptotic cell death, potentially with bystander effects on adjacent antigen-negative cells, as presented in Figure 1 [3, 4]. However, the net functional activity of an ADC is determined by the integrated action of its three components: the antibody, linker, and warhead, whose collective activity profile constitutes a complex, high-dimensional activity landscape. The activity landscape is challenging to explore experimentally, and high rates of attrition remain prevalent due to unforeseen toxicity, inadequate target engagement, or failure to elicit the desired therapeutic effect [5].

**Figure 1.**
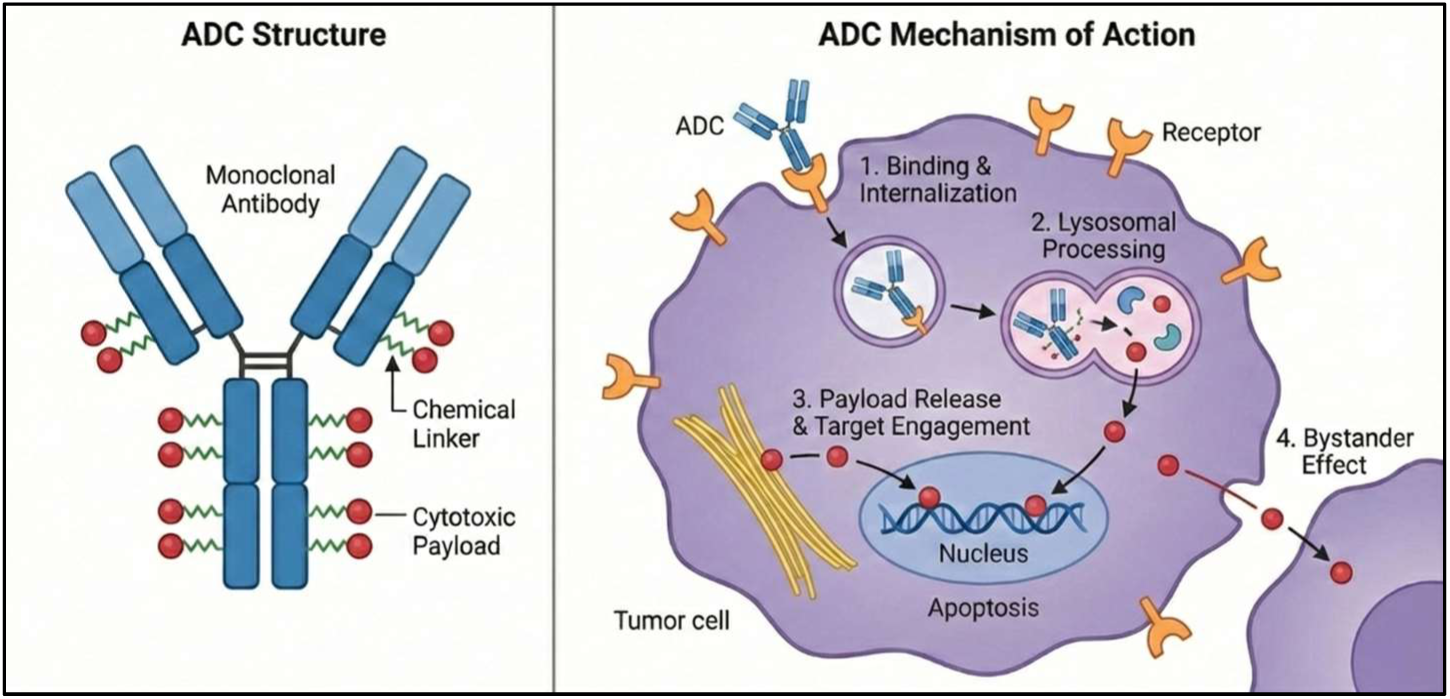
Structure and Mechanism of Antibody-Drug Conjugates. The left panel in Figure 1 shows the composition of the ADC, which consists of a monoclonal antibody conjugated to a cytotoxic payload via a linker and the right panel shows the mechanism of action of the ADC.

The increasing complexity of ADC component combinations has therefore motivated the adoption of deep learning–based computational methods, not to design ADCs directly, but to predict their activity and filter out unlikely candidates early in the discovery pipeline. Recent advances include de novo antibody generation using generative models and protein language model–based prediction of antibody–antigen or drug–target interactions [6–9]. While these approaches effectively model isolated components or pairwise interactions, a key gap remains in predicting the activity of ADCs as integrated constructs. Currently, ADCNet represents the only established deep learning framework developed explicitly for ADC activity prediction [10]. However, its core limitation lies in representing the antibody, antigen, linker, and payload as independent feature sets concatenated contextual integration. This omission restricts the model’s ability to leverage the antibody-antigen interface, the primary determinant of antigen engagement and a critical contributor to ADC activity.

To address this limitation while accounting for the scarcity of publicly available ADC data, we introduce ABFormer, a transformer-based framework for ADC activity prediction. ABFormer employs a transfer learning strategy by repurposing AntiBinder, a bidirectional-attention-based hybrid protein encoder designed for precise antibody-antigen interaction prediction, pretrained on the MET antigen dataset and used here as a frozen feature extractor [11].

This enables the capture of high-resolution antibody-antigen interaction embeddings without requiring large-scale retraining on the limited ADCdb dataset [12]. Latent space analysis confirms that these MET-derived embeddings encode transferable molecular recognition signals across diverse ADC targets [13]. These contextual embeddings are then fused with linker, payload, and DAR features in a multi-modal architecture that overcomes the limitations of naive feature concatenation [14].

ABFormer is designed as an early-stage computational screening system rather than a mechanistic ADC simulator. For any candidate ADC specification, the model outputs a probability reflecting its likelihood of belonging to the active class. These predictions support practical in-silico triage: researchers can rapidly evaluate antibodies against libraries of linker-payload combinations or vice versa, to deprioritize inactive constructs before synthesis [15]. This substantially accelerates ADC discovery by focusing experimental resources on the most promising candidates [16].

While ABFormer incorporates detailed antibody-antigen context, the linker and payload are still represented independently. A fully comprehensive activity-prediction model would ideally encode higher-order interactions among all ADC components. However, datasets such as ADCdb are too limited in chemical diversity to support such high-capacity interaction modelling [12, 17]. As explored further in the provided supplementary file, architectures that introduce explicit linker-payload interaction mechanisms such as bidirectional cross-attention modules tend to overfit the sparse combinatorial space of 82 linkers and 71 payloads, learning memorized pairs rather than transferable chemical principles [12]. These models show reduced specificity and less stable behaviour under rigorous leave-pair-out and external benchmarking, indicating that the additional chemical attention layers amplify noise rather than improving generalization. Under these constraints, prioritizing the antibody-antigen interface provides the most reliable and meaningful gains in ADC activity prediction [18]. Comparative benchmarks against ADCNet and several machine-learning baselines confirm that the contextual antibody-antigen embeddings in ABFormer substantially improve predictive performance [19].

## 2. Materials and Methods

### 2.1. Dataset and Preprocessing

The original dataset consisted of 435 ADCs. During the feature construction pipeline, six samples were identified as having antibody sequences that could not be processed by the AntiBinder module [20]. All analyses and data splits of ABFormer were therefore performed on the remaining 429 valid ADCs. ADCNet and fingerprint-based baselines were run on all 435 samples because they were able to process the entire set. To rigorously evaluate model generalization, we employ two complementary splitting strategies that address different aspects of ADC discovery [21]:

#### Random Split

The data is split into training (80%, n=343), validation (10%, n=43), and test sets (10%, n=43) through random sampling. This allows the same antibody/antigen to occur in each set with different linker payload combinations, simulating the exploration of known therapeutic targets with different conjugation chemistries [22].

#### Leave-Pair-Out Split

Each ADC is assigned a unique Pair ID based on exact sequence identity of its heavy chain, light chain, and antigen, yielding 149 distinct antibody-antigen pairs. Data are split at the pair level: training contains the corresponding unique pairs (340 ADCs), while validation and test contain unique pairs (43 and 46 ADCs, respectively). This ensures no antibody-antigen pair in training appears in validation or test sets, regardless of linker-payload composition, while maintaining a consistent label distribution (∼64% positive) across all subsets [18, 23].

#### Critical distinction

In random split, an ERBB2-targeting antibody (ERBB2 is a well-characterized gene-encoded antigen) with linker A may appear in training, while the same antibody with linker B appears in testing the model has seen the target during training. In leave-pair-out split, if an ERBB2-targeting pair is assigned to the test set, it never appears in training with any linker-payload combination. This rigorously evaluates generalization to completely novel therapeutic targets while allowing linker-payload chemistries to overlap across splits, consistent with real-world scenarios where established conjugation strategies are applied to new biological targets [24].

#### Note on sequence similarity

Due to the sparsity and presence of single occurrence antibodies and antigens in the ADCdb dataset, homology clustering was not feasible. The dataset has 149 distinct antibody and antigen pairs out of 9,536 possible pairs, with a sparsity of 98.4%. In such a scenario, homology clustering would result in the formation of micro-clusters that are too small to be split into train, validation, and test sets [25]. Hence, we assign Pair IDs based on the exact identity of the heavy chain, light chain, and antigen sequences. This ensures that no antibody and antigen pair appears across splits, thus avoiding memorization of a particular biological target while retaining sufficient samples for effective model training and testing [26]

### 2.2. Feature Representation

ABFormer integrates representations from three pre-trained models, each capturing distinct molecular modalities.

#### 2.2.1. Protein Sequence Encoding

The sequence encoding of the antigen and antibody light chains was performed using the ESM-2 protein language model (650M parameters) [27]. ESM-2 (650M) was preferred over the more recent ESM-3 models because it offers a stable and well-validated sequence encoder with reduced computational requirements [28]. The model architecture is designed to be optimal for single-sequence representation, which is more suitable for our sequence-centric tasks. Besides the PLM encodings, we also calculated amino acid composition (AAC) features for all antigen, light chain, and heavy chain sequences [29]. The antibody heavy chain sequences were also encoded using IgFold [30] to obtain structural embeddings, with region-specific indexing to clearly demarcate the complementarity-determining regions (CDRs) from framework residues in the variable domain [31]. The heavy chain regions are indexed according to the abnumber CDR definition (Chothia numbering): framework regions (FR) indexed as 1, and CDR1/2/3 regions indexed as 3/4/5 to facilitate region-aware attention in AntiBinder [32].

#### 2.2.2. Small Molecule Encoding

The SMILES strings of the linker and payload molecules are encoded with FG-BERT [33], a molecular transformer pre-trained on about 1.45 million molecules using functional-group-level masked language modeling, resulting in 256-dimensional embeddings for each molecule [33]. Moreover, we calculate 167-bit MACCS fingerprints for both the linker and the payload molecules to obtain complementary hand-engineered chemical features [34]. We employ 167-bit MACCS keys instead of Morgan fingerprints (2048-bit) to reduce the dimensionality [35]. With only 340 training examples, high-dimensional Morgan fingerprints would cause overfitting and heavy computational costs without any advantage [36].

We have re-implemented the original TensorFlow-based FG-BERT in PyTorch to guarantee that all model parameters are contained in a single PyTorch stack, allowing for efficient multi-GPU training with DistributedDataParallel [37]. The pre-trained weights have been transferred from the original checkpoint. FG-BERT is fine-tuned as a whole during the ABFormer training process.

#### 2.2.3. Antibody-Antigen Binding Interface Encoding

The AntiBinder is a bidirectional attention network pre-trained on the MET dataset (4,000 antibody-antigen pairs) for binding prediction [11]. AntiBinder combines sequence-level information from ESM-2 with structural information from IgFold via stacked cross-attention layers, resulting in 2592-dimensional interaction-aware embeddings [27, 30].

To predict ADC activity, we leverage the pre-trained AntiBinder as a frozen feature extractor, stripping away its classification head and taking the final hidden state as input to ABFormer. This transfer learning strategy allows us to model antibody-antigen interactions without extensive retraining on the small ADCdb dataset [12].

##### Domain generalization

AntiBinder was trained solely on MET (4,000 pairs), while ADCdb has 62 different antigens of cancer with no overlap with MET [38]. To ensure generalization, we conducted an extensive analysis of the latent space using UMAP dimensionality reduction, cluster quality estimation, and analysis of pairwise distance structures (will be discussed in Section 3.5) [39]. The results show high-quality separation of antigens (Silhouette = 0.685) and structured representation of the embeddings with intra-antigen distances 4.8 times smaller than inter-antigen distances (p < 0.001), thus proving that the learned representations generalize well from MET to other ADC antigens [40].

The pretrained model AntiBinder was trained on dataset that is completely separate from ADCdb, on the MET dataset. The dataset doesn’t include ADCs or experimental ADC-related data, thus preventing any information leakage from the pre-training step to the evaluation step.

### 2.3. ABFormer Architecture

#### 2.3.1 Model Overview

ABFormer is a multi-modal fusion model developed to predict the activity of ADC molecules by modeling the binding context of antibodies and antigens, as well as the payload, and conjugation properties between the payload and linker independently. The main idea of ABFormer is to use transfer learning to emphasize representations of the antibody-antigen interface, while also including the linker, payload, and Drug-Antibody Ratio (DAR) as shown in the structural overview in Figure 2. This is in contrast to previous models of ADC molecules, which were based on direct feature concatenation without emphasizing the binding information of the interfaces.

**Figure 2.**
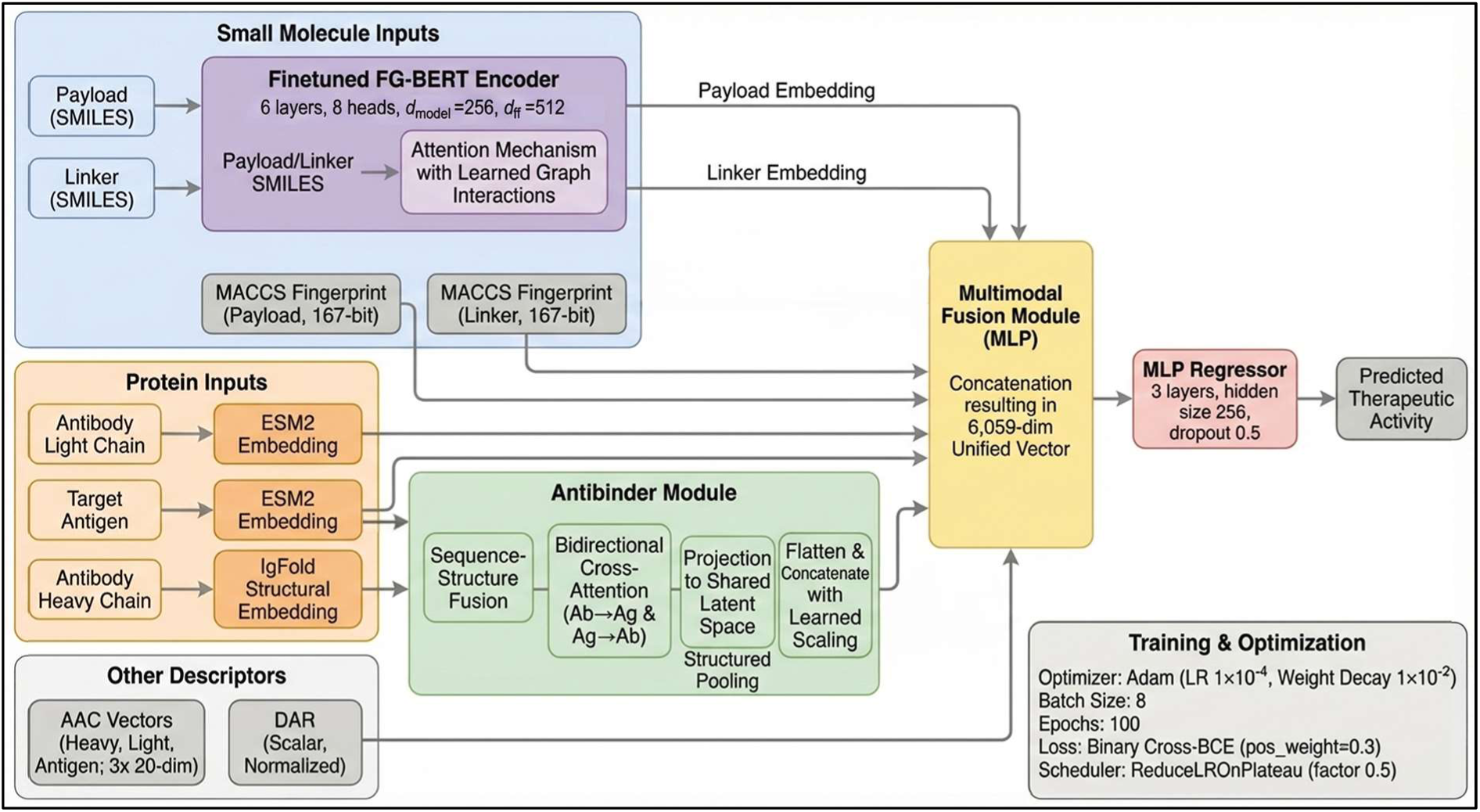
ABFormer’s Structural Overview. ABFormer’s structural overview in Figure 2 represents a multimodal ADC prediction framework that integrates drug features (DAR, payload, and linker), antigen sequences, and antibody sequences (heavy/light chains). Features are encoded via FG-BERT, ESM-2, Ig-Fold models, augmented with MACCS fingerprints and amino-acid composition (AAC) descriptors, and then fused through feature concatenation before passing through a fully connected network to generate the final prediction output.

ABFormer uses a pre-trained antibody-antigen interface encoder to capture transferable binding information, while fine-tuning the molecular encoders of the linker and payload to fit the chemical space of ADC molecules. The efficacy of this modular approach was confirmed by architectural ablation studies and fusion strategy comparisons discussed in provided supplementary file [41].

#### 2.3.2. Feature Integration and Fusion

Given an ADC sample, we extract the following feature representations:

- Antibody-antigen binding interface: 𝑡_1_ ∈ ℝ^2592 (from frozen AntiBinder)
- Antibody light chain: 𝑡_2_ ∈ ℝ^1280 (from ESM-2)
- Antigen sequence: 𝑡_3_ ∈ ℝ^1280 (from ESM-2)
- Linker: 𝑥_1_ ∈ ℝ^256 (from fine-tuned FG-BERT)
- Payload: 𝑥_2_ ∈ ℝ^256 (from fine-tuned FG-BERT)
- DAR: 𝑡_4_ ∈ ℝ^1 (normalized)
- Linker MACCS keys: 𝑥^MACCS^ ∈ ℝ^167^
- Payload MACCS keys: 𝑥^MACCS^ ∈ ℝ^167^
- Antibody Heavy Chain AAC featurization: 𝑎𝑎𝑐_1_ ∈ ℝ^20^
- Antibody Light Chain AAC featurization: 𝑎𝑎𝑐_2_ ∈ ℝ^20^
- Antigen AAC featurization: 𝑎𝑎𝑐_3_ ∈ ℝ^20^

These features are concatenated in the following order:

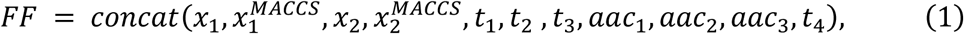

Resulting in a 6059-dimensional feature vector.

**Prediction Head**:

The concatenated feature vector is fed into a multi-layer perceptron (MLP) with GeLU activations, configured with layer dimensions 6059 → 256 → 256 → 1 [42]. ABFormer contains 4,784,641 trainable parameters, compared to ADCNet’s 4.38M, primarily allocated to accommodate the expanded dimensionality of AntiBinder embeddings (2592-dim vs. 1280-dim for independent heavy chain encoding) [10].

#### 2.3.3. Training Protocol Selective Fine-tuning Strategy

To balance transfer learning with task-specific specialization, the model is trained under a selective update strategy in which AntiBinder, ESM-2, and IgFold remain frozen and operate solely as feature extractors, while FG-BERT is fully fine-tuned and the MLP layers are trained from scratch. This setup enables FG-BERT to adapt to ADC-relevant chemical patterns despite the constrained linker–payload search space. The training setup uses the Adam optimizer (𝑙𝑟 = 1 × 10^-4^, 𝑤𝑑 = 1 × 10⁻²), a ReduceLROnPlateau scheduler (𝑚𝑜𝑑𝑒 = max, 𝑓𝑎𝑐𝑡𝑜𝑟 = 0.5, 𝑝𝑎𝑡𝑖𝑒𝑛𝑐𝑒 = 7), early stopping with a 30-epoch patience based on validation AUC, a batch size of 8, and a maximum of 100 epochs [37, 43]. The loss function is binary cross-entropy with logits, and all experiments run on a single NVIDIA A100-SXM4-40GB GPU, with training and testing repeated across random seeds 1–15 to ensure robustness and reproducibility.

#### 2.3.4. Architectural Design Justification

The choice to model antibody–antigen interactions explicitly and use independent linker and payload embeddings was justified through a comparison with six other architectural variants that are discussed in the provided supplementary file. Architectures that modeled linker–payload interactions explicitly through cross-attention and bilinear fusion modules or used alternative pre-trained molecular encoders such as MolFormer or MegaMolBART instead of FG-BERT generally showed suboptimal generalization performance [44, 45]. Although some of these variants were competitive during the optimization process, they performed worse on novel benchmark data, which suggested a lack of transferability.

Analysis of training dynamics showed that models with more modelling capacity on the linker–payload branch tended to overfit quickly, converging rapidly without learning representations that generalized well across ADCs. This was further worsened by the limited chemical diversity in the dataset, which has only 82 unique linkers, 71 unique payloads, and 340 training examples. In such a setting, modelling linker–payload interactions directly led to the memorization of sparse combinatorial patterns rather than learning robust chemical patterns [46].

Also, replacing FG-BERT with other molecular transformers led to deteriorated downstream performance, indicating that these molecular encoders failed to provide sufficiently informative and task-relevant functional group representations to add value in limited-data settings. On the contrary, FG-BERT’s chemistry-inspired embeddings always added value to the antibody-antigen representations without causing instability or overfitting [33].

Taken together, these findings suggest that the distribution of architectural complexity to the modeling of antibody–antigen interfaces via transfer learning is a far more effective strategy than learning linker–payload interactions from scratch. These findings support the notion that, within the current data paradigm, the primary factor in accurate ADC activity prediction is the transferable representation of antibody–antigen binding, with independent small-molecule encodings being sufficient [47].

### 2.4. Baseline Models and Training Details

#### 2.4.1. ADCNet Implementation

ADCNet serves as the primary baseline, utilizing its established architecture: ESM-2 for proteins (frozen), FG-BERT for small molecules (fine-tuned), and an MLP fusion layer. Crucially, all ADCNet results reported in this study were obtained by training the model from scratch on the same leave pair out data split, random split strategy and random seeds used for ABFormer. We did not rely on pre-trained weights or prior checkpoints, ensuring a strictly fair comparison under the same experimental conditions.

#### 2.4.2. Machine Learning Baselines

Four traditional ML models were evaluated: Logistic Regression (LR) [48], Random Forest (RF) [49], Support Vector Machine (SVM) [50], and XGBoost (XGB) [51].

##### Feature Representation

For small-molecule components (linker and payload), we computed either MACCS keys (167-bit) or Morgan fingerprints (2048-bit, radius=2) [52]. For protein components (antibody heavy chain, light chain, and antigen), sequences were encoded using

Amino Acid Composition (AAC), representing the normalized frequency of each of the 20 standard amino acids [53].

##### Feature Dimensionality

MACCS-based: 2 × 167 (molecules) + 3 × 20 (proteins) + 1 (DAR) = 395 features

Morgan-based: 2 × 2048 (molecules) + 3 × 20 (proteins) + 1 (DAR) = 4,157 features

We trained and tuned models independently on both the random split and the leave-pair-out split, each providing its own training, validation, and test partitions. Hyperparameter optimization was performed using the TPE algorithm (Hyperopt), maximizing ROC-AUC on the validation subset associated with whichever split and seed was being used [54, 55].

#### 2.4.3. Alternative Deep Learning Architectures

Six additional architectures incorporating linker-payload interaction modules or alternative encoders were evaluated. Detailed descriptions, training protocols, and results are in the provided supplementary file. These models were trained identically to ABFormer (same splits, seeds, hyperparameters) to isolate architectural effects.

### 2.5. Evaluation of model performance

The performance of the ABFormer model as well as that of the baseline ADCNet model is evaluated with a complete set of classification performance metrics that include positive predictive value (PPV), negative predictive value (NPV), specificity (SP/TNR), sensitivity (SE/TPR/Recall), accuracy (ACC), F1 score, balanced accuracy (BA), Matthews correlation coefficient (MCC)[56], area under the precision-recall curve (PRAUC) [23], and area under the receiver operating characteristic curve (AUC-ROC)[56–58]. The formulae for the metrics are defined as follows:

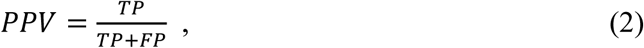

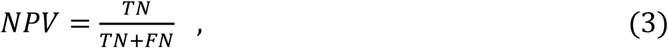

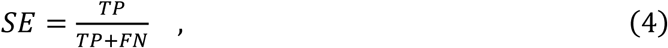

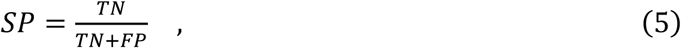

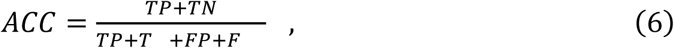

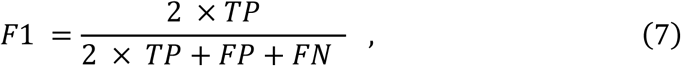

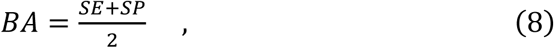

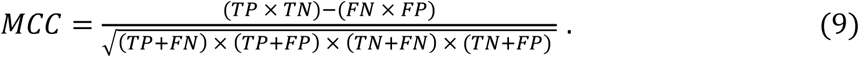

Where TP, TN, FP, FN, TNR, and TPR represent true positive, true negative, false positive, false negative, actual negative rate, and true positive rate, respectively. Furthermore, AUC-ROC is the area under the receiver operating characteristic curve plotted by calculating TPR vs. FPR at different discrimination thresholds, which measures the overall discriminative performance of the model. PRAUC represents the area under the precision-recall curve, which is particularly informative for evaluating performance on imbalanced datasets by plotting precision against recall at various threshold settings.

## 3. Results and discussion

### 3.1. ADC Modelling Dataset

The ABFormer model was trained on a curated subset of the ADCdb dataset. Following removal of six ADC entries whose antibody sequences could not be processed by the AntiBinder module due to invalidity or incompleteness, the final dataset consisted of 429 valid ADCs, comprising 144 unique antibodies, 62 antigens, 81 linkers, 70 payloads, and 74 distinct DAR values, yielding 144 unique antibody–antigen pairs. Activity labels assigned at 1000 nM produced 279 active and 150 inactive ADCs, reflecting moderate class imbalance. The associated chemical space exhibited broad diversity, with linker molecular weights ranging from 120.195-1702.11 Da and payloads from 233.097-1580.593 Da, and wide variation in AlogP distributions (linkers - 6.504 to 6.908; payloads - 4.675 to 7.406), indicating broad physicochemical coverage.

A detailed topology analysis of the dataset revealed significant structural sparsity and highly skewed antigen frequency distribution. Specifically, 69.8% of antibodies and 50% of antigens occurred exactly once in the dataset, while the most frequent antigen accounted for 25.5% of all ADCs and the top five antigens accounted for 57.2% of observations. Additionally, only 149 unique antibody–antigen pairs were represented across the entire 429-sample dataset, indicating substantial fragmentation [24].

To determine if homology-based clustering could help alleviate this sparsity problem, we analyzed the sequence similarity of all 62 distinct antigens. We calculated the pairwise sequence identity (percent similarity) with relaxed criteria (80%, 85%, 90%, 95%, 98%) to look for homologous antigen clusters [59]. Notably, all 62 antigens were found to be individual singleton clusters (groups of size 1) at all levels of sequence identity examined, with no antigen sequence showing ≥80% identity to any other antigen sequence. This result shows that the antigens are naturally distributed at the sequence level and cannot be clustered by homology.

Consequently, we used the exact sequence identity to assign a distinct Pair ID for each heavy chain, light chain, and antigen. This approach guarantees three essential aspects: (1) the absence of any antibody-antigen pair in both the training and test sets, thus preventing the model from memorizing particular targets; (2) the test antigens are true novel targets not observed in the training set; and (3) the overlap of linker and payload in the splits captures real-world settings where effective conjugation methods are employed for novel targets. The dataset was divided into training (n=340, 95 unique pairs), validation (n=43, 24 unique pairs), and test (n=46, 25 unique pairs) sets, with 58.8% of the test antigens not present in the training set.

The overlap of antigens between the training and test sets was moderate (58.8%), but the overlap at the antibody level was small (heavy chain 4.0%; light chain 19.0%), thus ensuring that the generalization performance is assessed mainly on novel antibody-antigen pairs, as expected in real-world ADC discovery. The label distribution for each split was balanced, and the Kolmogorov-Smirnov tests verified that there were no significant changes in the distribution of the subsets [60].

For the purpose of baseline assessments, the ADCNet and conventional machine learning models, which process explicit chemical fingerprints or other feature transformations, were assessed on all 435 original samples to ensure compatibility with their processing requirements.

### 3.2. Performance under Random Split (RS) Evaluation

We first evaluated all models using a basic Random-Split (RS) protocol, a common baseline approach in general deep learning experiments. Under this setting, several baseline methods, most notably ADCNet and traditional machine learning (ML) models achieved strong aggregate performance across sensitivity and accuracy metrics (Supplementary Table T1). In contrast, ABFormer did not exhibit a clear advantage under RS, particularly in terms of sensitivity, and in some cases appeared to underperform relative to ADCNet and ML baselines. However, closer inspection revealed that the apparent superiority of the baselines in RS was largely driven by aggressive positive predictions. High sensitivity was consistently accompanied by a pronounced drop in specificity (e.g., ADCNet 𝑆𝑃 = 0.6219), indicating a tendency to favor the majority “Active” class. This imbalance suggests that RS evaluation inflates performance by allowing models to exploit structural redundancy and scaffold overlap between training and test sets, rather than requiring genuine extrapolation to unseen antibody-antigen or linker-payload combinations [22].

ABFormer, by contrast, maintained a more balanced performance profile under RS, with substantially higher specificity (𝑆𝑃 = 0.8574), indicating a more conservative decision boundary. While this led to lower apparent sensitivity in RS, it suggested that ABFormer was less reliant on memorization and more focused on learning discriminative features.

### 3.3. Assessment via Leave-Pair-Out (LP) Validation

To more rigorously evaluate generalization, we next employed a Leave-Pair-Out (LP) cross-validation strategy, which explicitly removes antibody-antigen pairs from the training set and thus better reflects a cold-start ADC discovery scenario [61]. Under LP evaluation, ABFormer demonstrated a clear and consistent advantage over ADCNet across nearly all critical metrics as tabulated in Table 1, including specificity, Matthew’s correlation coefficient (MCC), and overall accuracy.

**Table 1.** Comparative Performance on Leave-Pair-Out (LP).

| Model | SE | SP | MCC | ACC | AUC | F1 | BA | PRAUC | PPV | NPV |
| --- | --- | --- | --- | --- | --- | --- | --- | --- | --- | --- |
| **ABFo**<br>**rmer** | 0.8023± | 0.9137± | 0.6964± | 0.8435± | 0.8983± | 0.8668± | 0.8580± | 0.9520± | 0.9475± | 0.7299± |
|  | 0.0243 | 0.1194 | 0.0895 | 0.0331 | 0.0061 | 0.0218 | 0.0505 | 0.0024 | 0.0700 | 0.0167 |
| **ADC**<br>**Net** | 0.6938± | 0.7519± | 0.4454± | 0.7170± | 0.8420± | 0.7397± | 0.7228± | 0.8734± | 0.8069± | 0.6385± |
|  | 0.1252 | 0.0476 | 0.0902 | 0.0596 | 0.0115 | 0.0779 | 0.0439 | 0.0132 | 0.0160 | 0.0889 |
| **SVM**<br>**(MAC**<br>**CS)** | 0.7644± | 0.7255± | 0.4852± | 0.7504± | 0.8210± | 0.7923± | 0.7450± | 0.9176± | 0.8283± | 0.6524± |
|  | 0.1377 | 0.1400 | 0.2503 | 0.1250 | 0.1080 | 0.1095 | 0.1241 | 0.0489 | 0.0855 | 0.1715 |
| **RF**<br>**(MAC**<br>**CS)** | 0.9022± | 0.7922± | 0.7004± | 0.8624± | 0.9391± | 0.8935± | 0.8472± | 0.9656± | 0.8856± | 0.8210± |
|  | 0.0235 | 0.0733 | 0.0759 | 0.0340 | 0.0226 | 0.0254 | 0.0416 | 0.0292 | 0.0369 | 0.0402 |
| **LR**<br>**(MAC**<br>**CS)** | 0.5822± | 0.6745± | 0.2482± | 0.6156± | 0.7059± | 0.6594± | 0.6284± | 0.8586± | 0.7642± | 0.4758± |
|  | 0.0213 | 0.0991 | 0.0850 | 0.0284 | 0.0181 | 0.0159 | 0.0430 | 0.0100 | 0.0570 | 0.0277 |
| **XGB**<br>**(Morgan)** | 0.8978±<br>0.0427 | 0.8275±<br>0.0608 | 0.7252±<br>0.0894 | 0.8723±<br>0.0426 | 0.9454±<br>0.0235 | 0.8996±<br>0.0347 | 0.8626±<br>0.0449 | 0.9730±<br>0.0118 | 0.9020±<br>0.0342 | 0.8233±<br>0.0634 |
| **SVM**<br>**(Morgan)** | 0.7422±<br>0.1087 | 0.6627±<br>0.1414 | 0.3987±<br>0.2434 | 0.7135±<br>0.1182 | 0.7993±<br>0.1039 | 0.7663±<br>0.0993 | 0.7025±<br>0.1226 | 0.8908±<br>0.0588 | 0.7930±<br>0.0901 | 0.5996±<br>0.1523 |
| **RF**<br>**(Morgan)** | 0.9422±<br>0.0235 | 0.7529±<br>0.0456 | 0.7238±<br>0.0421 | 0.8738±<br>0.0188 | 0.9316±<br>0.0180 | 0.9050±<br>0.0142 | 0.8476±<br>0.0224 | 0.9653±<br>0.0106 | 0.8711±<br>0.0204 | 0.8829±<br>0.0415 |
| **LR**<br>**(Morgan)** | 0.7556±<br>0.0272 | 0.7020±<br>0.0520 | 0.4477±<br>0.0412 | 0.7362±<br>0.0176 | 0.7708±<br>0.0133 | 0.7851±<br>0.0150 | 0.7288±<br>0.0221 | 0.8860±<br>0.0061 | 0.8183±<br>0.0244 | 0.6197±<br>0.0224 |

LP in table 1 is reported as mean ± standard deviation over 15 seeds for baseline ML models, ADCNet, and ABFormer.

While several ML baselines, particularly Random Forest and XGBoost using Morgan fingerprints achieved high sensitivity, this was again accompanied by reduced specificity, indicating recall-biased behavior. In contrast, ABFormer achieved the highest specificity and precision among all evaluated models, reflecting superior robustness against false positives. The resulting improvement in MCC highlights ABFormer’s ability to balance sensitivity and specificity under strict generalization constraints [56]. Importantly, the divergence between RS and LP results underscores a key limitation of random splitting: models that perform well under RS do not necessarily generalize to unseen antibody-antigen combinations. The LP results indicate that ABFormer’s architecture captures biologically meaningful interaction patterns rather than relying on shallow correlations [62].

### 3.4. Evaluation on Independent External Benchmark

To determine true real-world generalizability and eliminate the possibility of overfitting to such small dataset, we conducted a final evaluation on an independent benchmark of 22 ADCs. This benchmark includes 21 ADCs from the ADCNet study and one additional ADC sourced from an earlier patent, spanning multiple antibodies, antigens, linkers, payloads, and DAR values [63–68]. Similarity analysis revealed moderate-to-high structural dissimilarity (sequence-based for proteins, ECFP-4 for small molecules) relative to the training data, making the benchmark a stringent test of model robustness [52]. The complete benchmark is summarized in a Supplementary Table T5 that reports ADC identifiers, molecular components, similarity metrics, references, DAR values, and labels. Across 15 independent seeds, all baseline ML models and ADCNet exhibited a critical failure mode: despite their strong performance under RS and competitive metrics under LP, each assigned uniformly high predicted probabilities (>0.80) to nearly all benchmark samples, including confirmed negative controls (e.g., Sample 3 and Sample 22). This indicates severe miscalibration and confirms that these models behave as majority-class predictors when faced with previously unseen ADCs [69]. Their RS and LP success thus reflects memorization or class-frequency biases rather than transferable activity understanding.

In sharp contrast, ABFormer achieved **100% accuracy** on the benchmark. It consistently assigned low probabilities (P < 0.5) to both negative samples, while maintaining high confidence on all true positives. The clean separation between positive and negative predictions presented in Table 2 demonstrates that ABFormer has learned a genuinely generalizable representation of ADC bioactivity, successfully extrapolating across variations in antibodies, antigens, linkers, payloads, and DAR values. This benchmark evaluation provides the strongest evidence that ABFormer is the only model among those tested capable of reliable out-of-distribution generalization in ADC activity prediction [70].

**Table 2.** Mean predicted probability ± standard deviation.

| ADC Name | True Label | ABFormer | ADC Net | XGBoost (Morgan) | RF (Morgan) | SVM (Morgan) | LR (Morgan) | XGBoost (MACCS) | RF (MACCS) | SVM (MACCS) | LR (MACCS) |
| --- | --- | --- | --- | --- | --- | --- | --- | --- | --- | --- | --- |
| **H1D8-DC** | 1 | 0.8842<br>±<br>0.0158 | 0.9326<br>±<br>0.0769 | 0.9172<br>±<br>0.0518 | 0.8024<br>±<br>0.0375 | 0.9572<br>±<br>0.0264 | 0.9997<br>±<br>0.0004 | 0.9491<br>±<br>0.0217 | 0.7978<br>±<br>0.0271 | 0.9570<br>±<br>0.0223 | 0.9885<br>±<br>0.0071 |
| **Cetuximab-MAGNET-Gem** | 1 | 0.7517<br>±<br>0.0375 | 0.9976<br>±<br>0.0034 | 0.9371<br>±<br>0.0390 | 0.8419<br>±<br>0.0215 | 0.8269<br>±<br>0.0431 | 0.9497<br>±<br>0.0302 | 0.9474<br>±<br>0.0364 | 0.8391<br>±<br>0.0267 | 0.9028<br>±<br>0.0680 | 0.3824<br>±<br>0.3550 |
| **Trastuzumab-MAGNET-Gem** | 0 | **0.4790**<br>±<br>**0.0442** | **0.9978**<br>±<br>**0.0031** | **0.8939**<br>±<br>**0.0468** | **0.8327**<br>±<br>**0.0278** | **0.8134**<br>±<br>**0.0449** | **0.5195**<br>±<br>**0.1238** | **0.9211**<br>±<br>**0.0384** | **0.7905**<br>±<br>**0.0448** | **0.9025**<br>±<br>**0.0584** | **0.1630**<br>±<br>**0.2697** |
| **Sacituzumab-AE** | 1 | 0.6828<br>±<br>0.0494 | 0.9674<br>±<br>0.0503 | 0.8326<br>±<br>0.0947 | 0.8034<br>±<br>0.0228 | 0.7277<br>±<br>0.1250 | 0.2195<br>±<br>0.1259 | 0.9426<br>±<br>0.0225 | 0.7949<br>±<br>0.0211 | 0.8484<br>±<br>0.0399 | 0.4171<br>±<br>0.1994 |
| **Sacituzumab-DM1** | 1 | 0.6705<br>±<br>0.0524 | 0.9969<br>±<br>0.0042 | 0.8320<br>±<br>0.0779 | 0.6714<br>±<br>0.0568 | 0.7761<br>±<br>0.0365 | 0.8957<br>±<br>0.0232 | 0.9033<br>±<br>0.0538 | 0.7436<br>±<br>0.0285 | 0.7822<br>±<br>0.0299 | 0.8937<br>±<br>0.0297 |
| **Sacituzumab-DXd** | 1 | 0.6692<br>±<br>0.0494 | 0.9828<br>±<br>0.0154 | 0.8133<br>±<br>0.0720 | 0.7575<br>±<br>0.0615 | 0.6630<br>±<br>0.0549 | 0.4664<br>±<br>0.0800 | 0.8247<br>±<br>0.0700 | 0.7450<br>±<br>0.0323 | 0.8247<br>±<br>0.1334 | 0.9943<br>±<br>0.0031 |
| **ADC-C1** | 1 | 0.8833<br>±<br>0.0240 | 0.9322<br>±<br>0.0567 | 0.9395<br>±<br>0.0321 | 0.8977<br>±<br>0.0348 | 0.8995<br>±<br>0.0474 | 0.9865<br>±<br>0.0073 | 0.9327<br>±<br>0.0303 | 0.8678<br>±<br>0.0310 | 0.8712<br>±<br>0.0262 | 0.9954<br>±<br>0.0019 |
| **ADC-C3** | 1 | 0.8824<br>±<br>0.0245 | 0.9777<br>±<br>0.0152 | 0.9452<br>±<br>0.0284 | 0.8865<br>±<br>0.0227 | 0.8676<br>±<br>0.0314 | 0.9714<br>±<br>0.0063 | 0.9252<br>±<br>0.0267 | 0.8589<br>±<br>0.0244 | 0.7934<br>±<br>0.0530 | 0.8000<br>±<br>0.0645 |
| **ADC-C4** | 1 | 0.8825<br>±<br>0.0244 | 0.9846<br>±<br>0.0106 | 0.9453<br>±<br>0.0285 | 0.8886<br>±<br>0.0241 | 0.8790<br>±<br>0.0279 | 0.9871<br>±<br>0.0041 | 0.9252<br>±<br>0.0267 | 0.8589<br>±<br>0.0244 | 0.7934<br>±<br>0.0530 | 0.8000<br>±<br>0.0645 |
| **ADC-C5** | 1 | 0.8900<br>±<br>0.0234 | 0.9967<br>±<br>0.0043 | 0.9537<br>±<br>0.0235 | 0.8935<br>±<br>0.0222 | 0.8719<br>±<br>0.0295 | 0.9989<br>±<br>0.0007 | 0.9491<br>±<br>0.0189 | 0.8691<br>±<br>0.0215 | 0.8456<br>±<br>0.0307 | 0.9914<br>±<br>0.0027 |
| **ADC-C6** | 1 | 0.8901<br>±<br>0.0232 | 0.9969<br>±<br>0.0042 | 0.9538<br>±<br>0.0235 | 0.8953<br>±<br>0.0241 | 0.8829<br>±<br>0.0249 | 0.9995<br>±<br>0.0004 | 0.9491<br>±<br>0.0189 | 0.8691<br>±<br>0.0215 | 0.8456<br>±<br>0.0307 | 0.9914<br>±<br>0.0027 |
| **ADC-C7** | 1 | 0.8824<br>±<br>0.0245 | 0.9884<br>±<br>0.0076 | 0.9454<br>±<br>0.0293 | 0.8875<br>±<br>0.0228 | 0.8696<br>±<br>0.0282 | 0.9771<br>±<br>0.0050 | 0.9252<br>±<br>0.0267 | 0.8589<br>±<br>0.0244 | 0.7934<br>±<br>0.0530 | 0.8000<br>±<br>0.0645 |
| **ADC-C8** | 1 | 0.8825<br>±<br>0.0244 | 0.9972<br>±<br>0.0039 | 0.9626<br>±<br>0.0284 | 0.8866<br>±<br>0.0229 | 0.8573<br>±<br>0.0314 | 0.9834<br>±<br>0.0043 | 0.9269<br>±<br>0.0267 | 0.8578<br>±<br>0.0214 | 0.8026<br>±<br>0.0482 | 0.8157<br>±<br>0.0664 |
| **ADC-C9** | 1 | 0.8825<br>±<br>0.0244 | 0.9924<br>±<br>0.0052 | 0.9455<br>±<br>0.0293 | 0.8896<br>±<br>0.0240 | 0.8804<br>±<br>0.0265 | 0.9897<br>±<br>0.0032 | 0.9252<br>±<br>0.0267 | 0.8589<br>±<br>0.0244 | 0.7934<br>±<br>0.0530 | 0.8000<br>±<br>0.0645 |
| **ADC-C10** | 1 | 0.8856<br>±<br>0.0234 | 0.9266<br>±<br>0.0571 | 0.9488<br>±<br>0.0287 | 0.8947<br>±<br>0.0282 | 0.8800<br>±<br>0.0429 | 0.9538<br>±<br>0.0088 | 0.9327<br>±<br>0.0325 | 0.8615<br>±<br>0.0267 | 0.8521<br>±<br>0.0413 | 0.9939<br>±<br>0.0017 |
| **ADC-C11** | 1 | 0.8954<br>±<br>0.0219 | 0.9984<br>±<br>0.0006 | 0.9688<br>±<br>0.0224 | 0.9109<br>±<br>0.0175 | 0.8835<br>±<br>0.0437 | 0.9981<br>±<br>0.0011 | 0.9534<br>±<br>0.0222 | 0.8779<br>±<br>0.0225 | 0.8626<br>±<br>0.0403 | 0.9986<br>±<br>0.0006 |
| **ADC-C12** | 1 | 0.8901<br>±<br>0.0233 | 0.9986<br>±<br>0.0005 | 0.9667<br>±<br>0.0222 | 0.9075<br>±<br>0.0179 | 0.8762<br>±<br>0.0253 | 0.9993<br>±<br>0.0004 | 0.9524<br>±<br>0.0163 | 0.8691<br>±<br>0.0207 | 0.8523<br>±<br>0.0317 | 0.9923<br>±<br>0.0023 |
| **ADC-C13** | 1 | 0.8950<br>±<br>0.0217 | 0.9985<br>±<br>0.0006 | 0.9674<br>±<br>0.0218 | 0.9120<br>±<br>0.0202 | 0.8972<br>±<br>0.0342 | 0.9995<br>±<br>0.0003 | 0.9531<br>±<br>0.0216 | 0.8807<br>±<br>0.0233 | 0.8552<br>±<br>0.0390 | 0.9943<br>±<br>0.0020 |
| **ADC-C14** | 1 | 0.8854<br>±<br>0.0230 | 0.9020<br>±<br>0.0831 | 0.9463<br>±<br>0.0280 | 0.8932<br>±<br>0.0285 | 0.8825<br>±<br>0.0402 | 0.9734<br>±<br>0.0088 | 0.9252<br>±<br>0.0336 | 0.8641<br>±<br>0.0298 | 0.7739<br>±<br>0.0735 | 0.7115<br>±<br>0.0691 |
| **ADCT-602** | 1 | 0.9355<br>±<br>0.0144 | 0.9739<br>±<br>0.0429 | 0.9523<br>±<br>0.0531 | 0.8449<br>±<br>0.0239 | 0.6811<br>±<br>0.1591 | 0.8195<br>±<br>0.0667 | 0.9410<br>±<br>0.0693 | 0.8573<br>±<br>0.0204 | 0.7815<br>±<br>0.1195 | 0.7257<br>±<br>0.1064 |
| **FZ-AD005** | 1 | 0.9827<br>±<br>0.0048 | 0.0437<br>±<br>0.0538 | 0.7597<br>±<br>0.1130 | 0.7507<br>±<br>0.0344 | 0.8294<br>±<br>0.0402 | 0.9942<br>±<br>0.0038 | 0.4133<br>±<br>0.1840 | 0.5432<br>±<br>0.0324 | 0.6415<br>±<br>0.1121 | 0.0096<br>±<br>0.0185 |
| CAT-01-106 | 0 | 0.4485<br>±<br>0.0466 | 0.8814<br>±<br>0.1184 | 0.9133<br>±<br>0.0505 | 0.8612<br>±<br>0.0296 | 0.8242<br>±<br>0.0619 | 0.9858<br>±<br>0.0061 | 0.9362<br>±<br>0.0623 | 0.8248<br>±<br>0.0198 | 0.8947<br>±<br>0.0883 | 1.0<br>±<br>0.0 |

Table 2 shows mean predicted probability ± standard deviation on the independent benchmark using 15-seed inference. Bold values indicate negative samples (<0.5).

### 3.5. Model Architecture Validation via Ablation Experiments

To delineate the specific contributions of the multi-modal components within ABFormer, we conducted a systematic ablation study. We evaluated model variants by selectively modifying the feature fusion vector 𝐹𝐹, originally defined in Eq. (1), while keeping the remaining architecture and training hyperparameters fixed.

The baseline feature fusion vector for the full model is defined as:

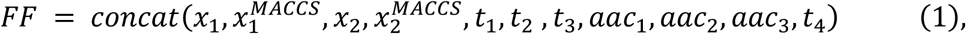

The resulting performance metrics, summarized Supplementary Table T4 and visualized in Figure 3, reveal that the contextual antibody-antigen interaction features are the primary driver of predictive success, while chemical embeddings are critical for minimizing false positives [71].

**Figure 3.**
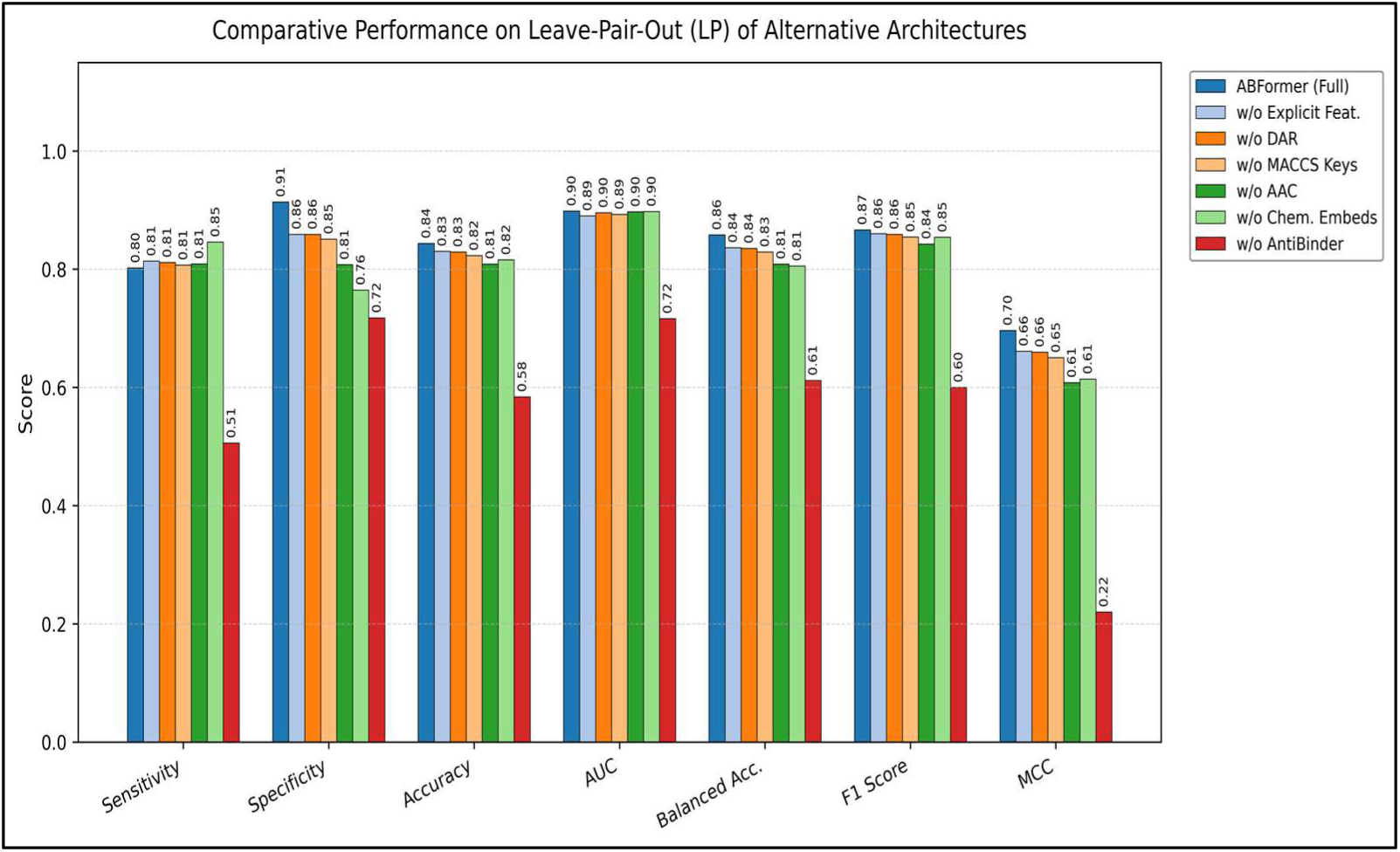
Comparative Performance of ABFormer Ablation Variants under Leave-Pair-Out Evaluation. The bar groups in Figure 3 summarize how the full ABFormer model performs relative to ablated variants where specific feature components are removed. Each color corresponds to a different architecture configuration (e.g., removal of DAR, MACCS keys, AAC, or chemical embeddings). Higher bars indicate stronger predictive performance across metrics such as sensitivity, specificity, accuracy, AUC, balanced accuracy, F1 score, and MCC. This legend clarifies which architectural element each variant lacks, allowing direct comparison of the contribution of each component to overall model robustness.

#### 3.5.1. The Critical Role of Contextual Antibody-Antigen Embeddings

The central hypothesis of this study is that explicit modelling of the antibody-antigen binding interface is required to accurately forecast ADC activity. We tested this by zeroing out the transfer-learning features derived from the pre-trained AntiBinder module 𝑡_1_(w/o AntiBinder), we did not remove the feature vector but replaced it with a zero tensor of identical dimensionality (0 ∈ ℝ^2592^ ) to strictly isolate the contribution of the learned signal without altering the MLP capacity, as shown in Eq. (10):

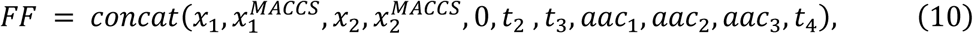

As shown in Figure 3, the Matthews Correlation Coefficient (MCC) dropped drastically from 0.6964 (Full Model) to 0.2199, and the classification accuracy plummeted to 58.41%, which is only slightly better than random chance. This is expected, since the independent component features alone (such as the antibody and antigen sequences) are not sufficient to capture the intricate activity landscape of ADCs. The AntiBinder module, which employs bi-cross attention to model the interplay between the heavy chain and the antigen, provides the necessary biological context that grounds the predictions of the model [11].

#### 3.5.2. Contribution of Chemical Encoders to Specificity

We further analysed the role of the fine-tuned FG-BERT chemical encoders. In this experiment (w/o Chemical Embeddings), the dense embeddings for the linker 𝑥_1_ and payload 𝑥_2_ were explicitly removed from the concatenation operation, forcing the model to rely solely on MACCS fingerprints for chemical context, as shown in Eq. (11):

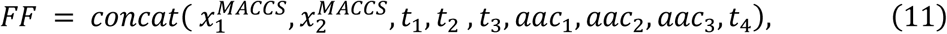

This modification produced a distinct failure mode characterized by high sensitivity (0.8460) coupled with substantially reduced specificity (0.7647). The resulting distortion of the performance polygon is clearly visible in Figure 3, where the sensitivity axis expands while the specificity axis contracts. This behaviour indicates that, in the absence of detailed chemical representations, the model becomes over-optimistic frequently predicting activity based on antibody targeting alone while failing to penalize chemically unfavourable linker-payload configurations. Thus, while biological context drives overall classification capability, the FG-BERT chemical encoders serve as a critical specificity filter.

#### 3.5.3. Impact of Explicit Feature Engineering

The removal of explicit features, specifically MACCS fingerprints (Eq. 12), Amino Acid Composition (AAC) (Eq. 13), and Drug-Antibody Ratio (DAR) resulted in minor performance decrements, as shown in Figure 3. The ablation of MACCS keys (w/o MACCS Keys) involved removing the explicit fingerprint vectors:

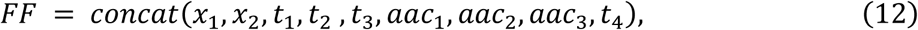

Similarly, for the ablation of AAC features (w/o AAC), the composition vectors were excluded:

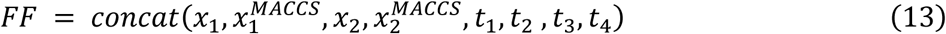

Removing MACCS keys reduced the MCC by approximately 0.046 (0.6964 to 0.6505). While less critical than the deep embeddings, these features provide incremental stability, likely by offering the downstream MLP direct access to fundamental physicochemical properties that reinforce the high-dimensional transformer representations [72].

### 3.6. Transfer Learning from MET to Diverse Cancer Antigens

AntiBinder was pre-trained on MET receptor binding data, yet ADCdb contains 429 samples spanning 62 unique cancer antigens with no MET representation. This raises a critical question: can embeddings learned from MET generalize to structurally diverse targets such as ERBB2 (Erb-B2 Receptor Tyrosine Kinase 2), KIT, FGFR2 (Fibroblast growth factor receptor 2), TNFRSF1A (Tumor Necrosis Factor Receptor Superfamily Member 1A), and EGFR (epidermal Growth Factor Receptor)? We performed latent space analyses to assess whether AntiBinder captured transferable binding features versus MET-specific patterns.

#### 3.6.1 Cluster Separation and Embedding Space Structure

We analysed embeddings for the five most prevalent antigens: ERBB2 (n=111), KIT (n=48), FGFR2 (n=36), TNFRSF1A (n=32), and EGFR (n=21), totalling 248 samples representing diverse protein families. UMAP projection (𝑛_𝑛𝑒𝑖𝑔ℎ𝑏𝑜𝑟𝑠 = 15, 𝑚𝑖𝑛_𝑑𝑖𝑠𝑡 = 0.1, 𝑚𝑒𝑡𝑟𝑖𝑐 = ′𝑐𝑜𝑠𝑖𝑛𝑒′) revealed five distinct, spatially separated clusters corresponding precisely to antigen targets as depicted in Figure 4 [39]. Quantitative assessment confirmed strong cluster quality:

**Figure 4.**
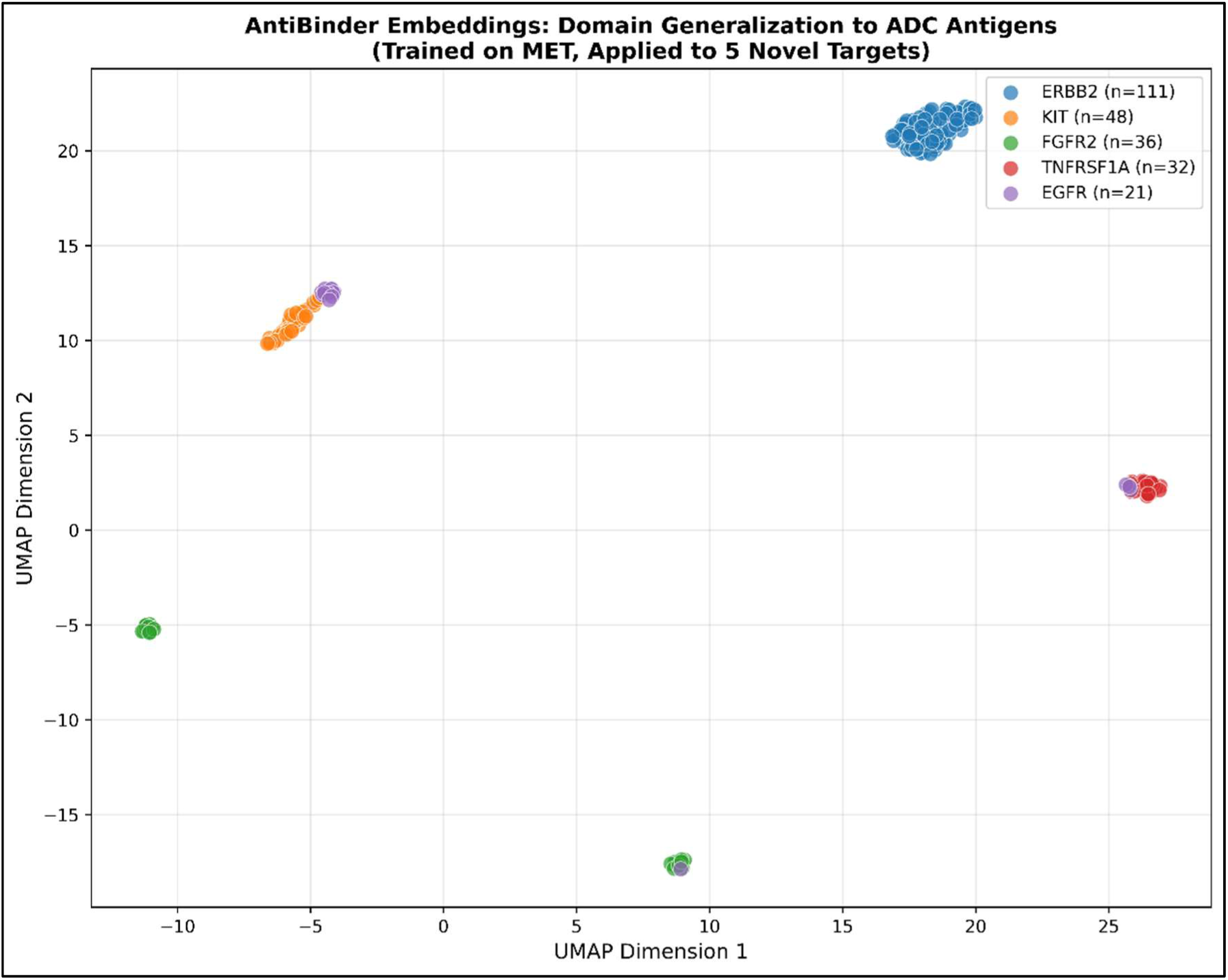
Visualisation of domain generalization of AntiBinder embeddings to diverse ADC antigens. Figure 4 presents UMAP projection of AntiBinder embeddings for 248 ADC samples across five antigen targets (ERBB2, KIT, FGFR2, TNFRSF1A, EGFR). Despite being trained exclusively on MET data, the model organizes samples into five distinct and well-separated clusters (Silhouette score = 0.710), indicating strong transfer of embedding-space structure to unseen antigen classes.

Silhouette score = 0.685 in high-dimensional space and 0.710 in 2D projection (values >0.5 indicate well-defined clusters). The Davies–Bouldin Index (DBI) for this clustering is 1.153 [73]. Because lower DBI values indicate more compact and well-separated clusters, this score suggests a reasonably good level of separation compared with typical clustering results.

To quantify embedding space geometry, we computed all pairwise cosine distances (30,628 pairs) and categorized them as intra-antigen (same target, n=8,569) or inter-antigen (different targets, n=22,059). Intra-antigen distances showed mean = 0.056 ± 0.083, while inter-antigen distances showed mean = 0.265 ± 0.049, yielding a 4.8-fold ratio with substantially separated distributions as shown in Figure 5. Statistical testing confirmed highly significant separation between intra- and inter-antigen distances (Mann–Whitney U test, p < 0.001) [74], with a large absolute gap in cosine distance distributions (Figure 5).

**Figure 5.**
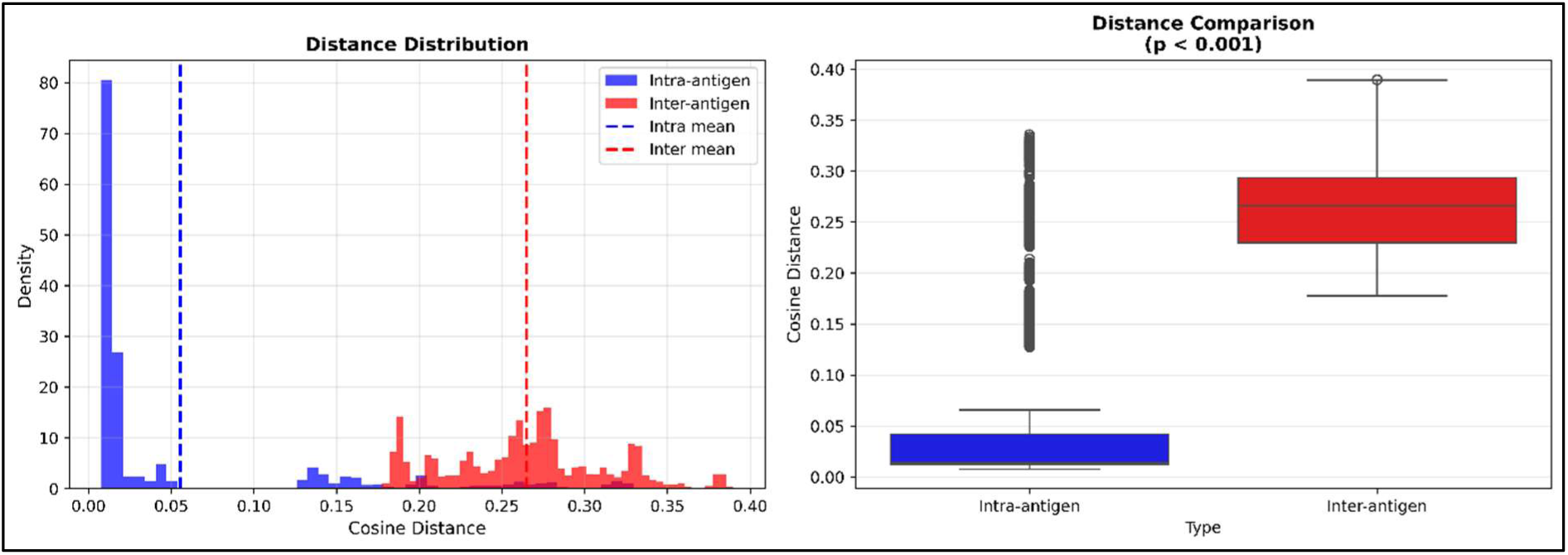
Quantitative analysis of embedding space structure. Figure 5 presents distribution of pairwise cosine distances (left) in the 2,592-dimensional embedding space. Blue: intra-antigen distances (same target, mean = 0.056). Red: inter-antigen distances (different targets, mean = 0.265). Dashed lines denote group means and box-plot comparison (right) showing a 4.8-fold difference between intra- and inter-antigen distances (Mann–Whitney U test: p < 0.001), confirming robust antigen-specific structure in the embedding space.

#### 3.6.2 Biological Interpretation

The successful generalization from MET to diverse ADC antigens reflects several contributing factors:

##### Shared Structural Features

Four of the five top antigens (ERBB2, KIT, FGFR2, EGFR) are Receptor Tyrosine Kinases (RTK) with modular extracellular domain architectures and conserved signalling paradigms. While their extracellular domains differ in detail, partial structural homology and shared modes of ligand and antibody engagement likely facilitate transfer of learned binding interface patterns across the RTK family [75].

##### Universal Binding Principles

Antibody-antigen recognition relies on general biophysical principles like shape complementarity, electrostatic interactions, hydrogen bonding that are largely antigen-agnostic. The bi-cross attention mechanism appears to capture these invariant features rather than MET-specific patterns [76].

##### Antibody Sequence Homology

A potential confounding factor is antibody similarity within antigen groups. For example, ERBB2-targeting ADCs include multiple trastuzumab derivatives with highly similar heavy chain sequences, which may contribute to intra-antigen clustering. While this indicates that part of the observed structure may reflect antibody sequence homology rather than binding interface physics alone, it does not contradict the generalization result. Successfully associating related antibody architectures with their corresponding antigens still requires transfer of learned representations from MET pretraining to unseen targets. The structured embedding space likely reflects a combination of sequence similarity and shared binding principles.

#### 3.6.3 Implications and Limitations

While our analysis provides strong evidence of transferable structure in the learned embedding space (Silhouette = 0.685, 4.8-fold intra- vs. inter-antigen distance ratio, p < 0.001), several limitations should be noted. The five most prevalent antigens analyzed are enriched for RTKs with partial structural similarity to MET; generalization to more divergent antigen classes such as integrins, GPCRs, or cytokines remains less well validated. In addition, clustering within antigen groups may partially reflect antibody sequence family effects rather than binding interface physics alone, although this still indicates successful transfer of learned representations from MET to unseen antigen contexts. Finally, generalization is assessed through embedding-space structure and clustering metrics rather than direct binding affinity prediction on held-out antigens.

Nevertheless, the combination of strong cluster separation, a highly structured embedding space, and consistent discrimination of five biologically distinct targets supports that MET-trained AntiBinder embeddings capture transferable binding context relevant for ADC activity prediction across diverse cancer antigens. These results suggest that bi-cross attention models trained on limited antibody-antigen data can learn molecular recognition patterns that extend beyond their original training distribution.

### 3.7. Latent Space Analysis and Validation of Learned Molecular Representations

To assess the feature learning behaviour of the ABFormer framework, we visualized the high-dimensional latent representations produced by the prediction subnetwork using t-distributed Stochastic Neighbour Embedding (t-SNE) [77]. The full dataset (training, validation, and testing samples) was used to project the learned multi-modal embeddings integrating antibody, antigen, linker, payload, and DAR features into two dimensions for qualitative comparison between untrained and trained models.

As shown in Figure 6 (Before Training), the untrained embedding space exhibits substantial overlap between positive (green) and negative (orange) samples. While limited local structure is visible, likely reflecting intrinsic molecular similarities, the two classes remain largely intermingled, indicating that the raw input representations do not provide clear class-discriminative structure. After training, however, the embedding space undergoes a pronounced reorganization. The trained ABFormer model produces a clear separation between positive and negative samples, forming distinct clusters in the low-dimensional projection. This qualitative separation is consistent with the model’s strong predictive performance, reflected by an AUC of 0.921 on the true labels. Together, these results suggest that training enables ABFormer to extract and integrate informative, non-linear features relevant to ADC activity [78].

**Figure 6.**
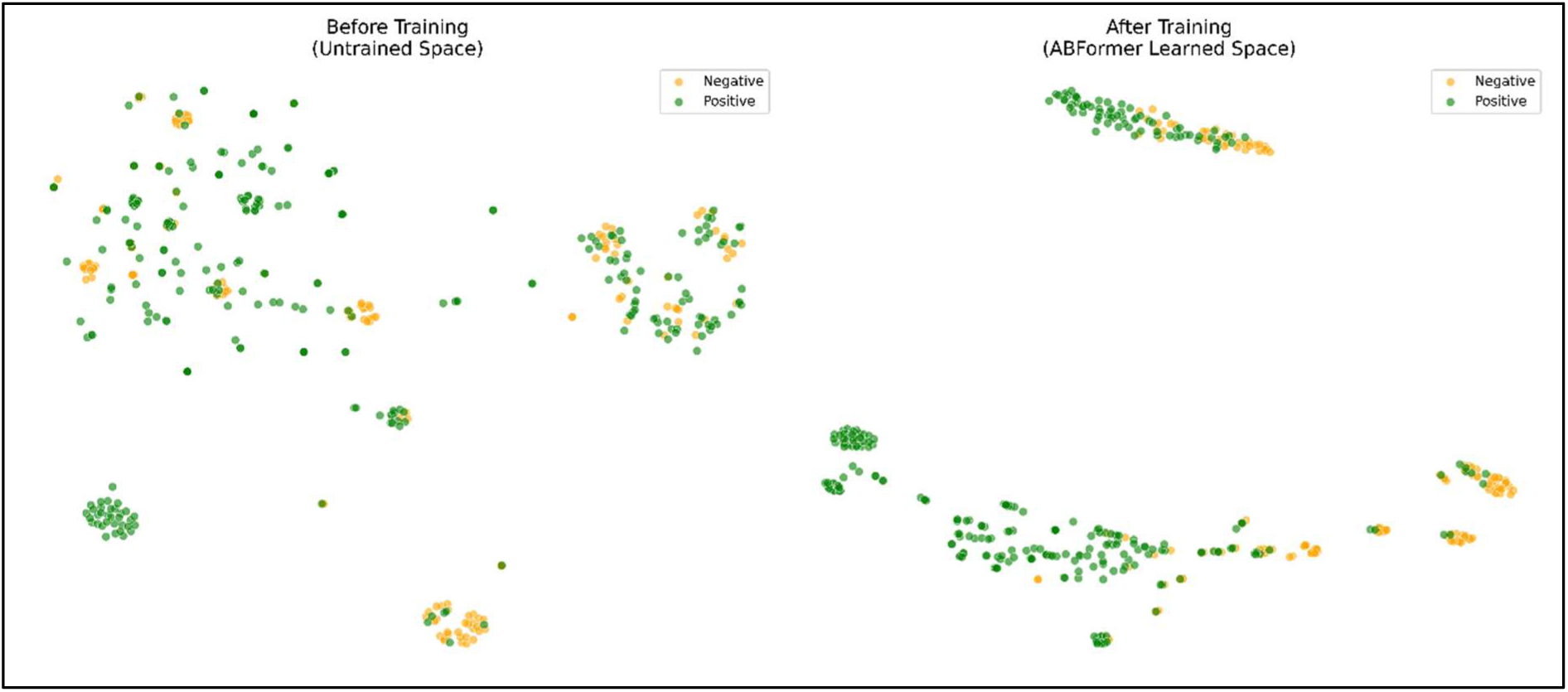
t-SNE visualization of ABFormer’s latent representation space before and after training. The plots in Figure 6 show 2-dimensional t-SNE projections of multi-modal ADC embeddings integrating antibody, antigen, linker, payload, and DAR features. **Left:** Untrained model (randomly initialized weights) displays substantial overlap between positive (green) and negative (orange) samples, indicating that raw input features lack intrinsic class-discriminative structure. **Right:** After training, ABFormer produces a distinctly organized latent space in which positive and negative samples separate into well-formed clusters. This emergent structure reflects the model’s ability to learn meaningful, non-linear biological and chemical relationships relevant to ADC activity.

To verify that the observed structure arises from genuine molecular feature learning rather than spurious correlations or overfitting, we performed a Y-scrambling validation as a negative control [79]. In this experiment, sample labels were randomly permuted while preserving all input features, thereby eliminating any true structure–activity relationships. As shown in Figure 7, the model trained on scrambled labels fails to produce meaningful class separation, yielding an embedding distribution resembling the untrained state with fully intermixed positive and negative samples. Correspondingly, predictive performance collapses from an AUC of 0.921 with true labels to 0.473 under Y-scrambling, which is indistinguishable from random guessing. The contrast between strong performance on genuine data and near-random behaviour under label permutation supports that ABFormer learns chemically meaningful patterns rather than memorizing dataset artifacts [70].

**Figure 7.**
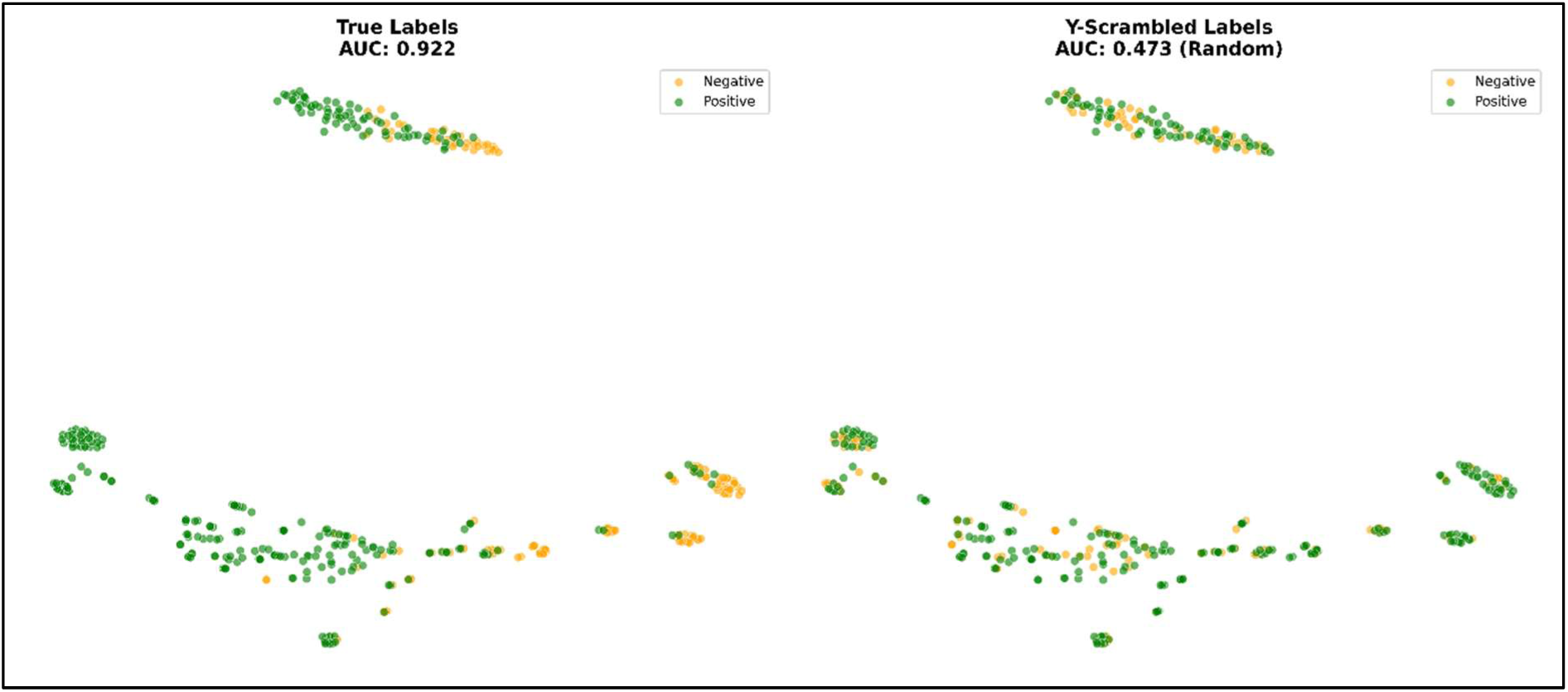
Y-scrambling validation of ABFormer’s learned latent space.

t-SNE projections in Figure 7 illustrate the effect of training under true versus randomly permuted labels. **Left:** With true activity labels (AUC = 0.922), ABFormer produces a structured latent space in which positive (green) and negative (orange) samples form well-separated clusters, reflecting meaningful molecular feature learning. **Right:** Under Y-scrambled labels (AUC = 0.473), the learned embedding space collapses into an intermixed distribution with no class separation, resembling an untrained or randomly organized space. The near-random AUC and loss of structure confirm that ABFormer’s predictive behaviour arises from genuine biological and chemical signal rather than memorization or spurious correlations.

## 4. Conclusion

In this study, we proposed ABFormer, a transformer-aided multi-modal model for predicting the activity of antibody-drug conjugates (ADCs) that captures the context of antibody and antigen binding through transfer learning. By leveraging a pre-trained AntiBinder model to generate interaction-aware antibody and antigen embeddings and combining them with chemically informed linker, payload, and DAR embeddings, ABFormer goes beyond the current state-of-the-art ADC modeling efforts, which are mostly based on independent feature concatenation.

Our results show that ABFormer generalizes significantly better than ADCNet and a variety of traditional machine learning baselines in rigorous generalization scenarios. Although random split evaluations expose the shortcomings of the baselines due to memorization and class bias, leave-pair-out validation and an external test set emphasize the generalization capabilities of ABFormer to novel antibody and antigen pairs and ADC molecules. In particular, ABFormer is the only model that correctly predicts the negative controls in true out-of-distribution tests, while all other models fail. Ablation studies further support that the main contribution of contextual antibody and antigen embeddings is predictive performance, while fine-tuned chemical encoders are essential for maintaining specificity and reducing false positives.

Latent space analyses and Y-scrambling experiments further support that ABFormer learns biologically and chemically meaningful representations, rather than relying on spurious correlations. Despite being trained in highly constrained data regimes, the model successfully transfers binding knowledge from a single-antigen pre-training setting (MET) to a variety of cancer targets, which highlights the effectiveness of interaction-centric transfer learning in low-data biomedical applications.

In conclusion, ABFormer opens a new avenue for ADC activity prediction, as it clearly shows that the priority of modeling the antibody and antigen interface is highly effective for improving robustness and real-world applicability. As larger and more diverse ADC datasets become available, this model can be naturally extended to incorporate higher-order interactions among all ADC components, further advancing the field of data-driven ADC design and virtual screening.

## List of abbreviations

ACC: Accuracy
ADC: Antibody Drug Conjugates
ADCdb: Antibody Drug Conjugates databank
AUC: Area Under Curve
BA: Balanced Accuracy
DAR: Drug-to-Antibody Ratio
ESM2: Evolutionary Scale Modeling version 2
Fc: Fragment Crystallisable
FFN: Feed Forward Network
FG-BERT: Fine Grained - Bidirectional Encoder Representations from Transformers
FN: False Negative
FP: False Positive
HER2: Human Epidermal Growth Factor Receptor 2
LR: Logistic Regression
MCC: Matthews Correlation Coefficient
MET dataset: Mesenchymal-Epithelial transition gene dataset
ML: Machine Learning
MLP: Multi Layer Perceptron
NPV: Negative Predictive Value
PPV: Positive Predictive Value
PRAUC: Precision-Recall AUC
QLoRA: Quantized Low-Rank Adaptation
RF: Random Forest
SE: Sensitivity
SP: Specificity
SVM: Support Vector Machine
TAA: Tumor Associated Antigen
TN: True Negatives
TNR: True Negative Rate
TP: True Positives
TPR: True Positive Rate
XGB: XG Boost

## Declarations

### Ethics approval and consent to participate

Not applicable

## Consent for publication

Not applicable

## Supplementary Material

Supplementary information and tables are provided as Supplementary File S1.

## Data availability

ADCdb data used in this study is publicly available from https://adcdb.idrblab.net/ All experiments were run on PyTorch [2.8.0. version] with CUDA [12.8 version]. **Code availability**

The code is available at GitHub Link: https://github.com/drugparadigm/ABFormer

## Author Contributions

**R.K.:** Investigation; validation; methodology; visualization; writing—original draft; formal analysis. **V.L.:** Data curation; formal analysis; methodology; validation; visualization. **S.G.:** Conceptualization; methodology; validation; visualization; formal analysis; data curation; software; supervision; formal analysis; writing—review and editing. **V.K.:** Conceptualization; methodology; writing—review and editing; supervision; formal analysis.

## Acknowledgements

We authors, Rushi Katabathuni, Viren Loka, Sanjana Gogte, and Vani Kondaparthi, express our sincere gratitude to the Drugparadigm Research Lab for providing the necessary facilities and infrastructure that enabled the successful completion of this work.

