## Supplementary File S1 for "ABFormer: A Transformer-based Model to Enhance Antibody-Drug Conjugates Activity Prediction through Contextualized Antibody-Antigen Embedding"

**Supplementary S1.**

**1. BCAA: Bilateral Cross-Attention Fusion**

**1.1. Architecture and Design**

The **BCAA** model extends ABFormer by retaining its AntiBinder-based biological backbone while replacing ABFormer’s independent chemical encodings with an explicit linker–payload interaction module. This module employs bidirectional cross-attention, allowing the linker to attend to the payload and vice versa, thereby producing a unified chemical interaction embedding intended to capture steric and stability-related effects. Incorporating this mechanism increases the parameter count to **5.3M** (vs. **4.7M** in ABFormer), with the additional ~600k parameters originating entirely from the chemical attention layers. The design is motivated by the assumption that ABFormer’s concatenation strategy oversimplifies the physics of the linker–payload bond.

**1.2. Training Dynamics (Seed 1)**

Both models converge, yet ABFormer achieves a higher validation peak (**AUC 0.8416**) than BCAA (**AUC 0.8258**). The reduced generalization of BCAA indicates that the expanded chemical module tends to memorize specific linker–payload combinations rather than learning reusable chemical principles.

**1.3. Leave-Pair-Out (LP) Performance (15 Seeds)**

Under LP evaluation, BCAA shows consistent regression relative to ABFormer. Specificity declines from **0.914 ± 0.12** (ABFormer) to **0.820 ± 0.14**, and MCC decreases from **0.696 ± 0.09** to **0.621 ± 0.09**. Accuracy and AUC also drop modestly. The specificity reduction reflects an increased false-positive rate, suggesting that the cross-attention layers occasionally overemphasize potent payloads independent of antibody context.

**1.4. Benchmark Analysis**

Independent benchmarking confirms that BCAA remains functional and does not collapse on negative controls. However, its outputs are consistently closer to the decision boundary and show higher variance (e.g., 0.48 ± 0.08 vs. ABFormer’s 0.45 ± 0.04). While the AntiBinder module reliably transmits the negative biological signal, the chemical cross-attention introduces uncertainty, making predictions less stable than those of ABFormer.

**1.5. Root Cause and Outlook**

The performance gap reflects a **data sufficiency limitation**. Learning cross-attentive linker–payload interactions requires a dense combinatorial dataset, yet ADCdb contains only ~80 linkers and ~70 payloads—insufficient for training the ~600k parameters of BCAA’s chemical module. Consequently, the model learns noise or memorized pairs, weakening the biological signal. ABFormer’s simpler chemical assumptions are more compatible with the current dataset scale. Nonetheless, BCAA remains a future-ready architecture: with a 5–10× expansion of ADC data, its higher capacity would likely enable it to surpass ABFormer by capturing interaction physics that simpler models cannot.

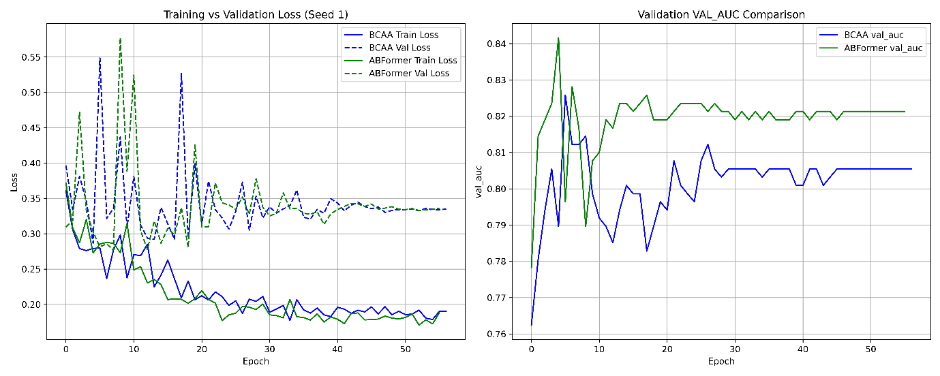

**Supplementary Figure 1.** Training Dynamics and Validation Performance Comparison Between BCAA and ABFormer. The left panel plots the train and validation loss curves for both models, with blue and green lines representing BCAA training and validation loss, and ABFormer training and validation loss, respectively. The right panel plots the validation AUC curves for both models, with blue and green lines representing BCAA and ABFormer, respectively.

**2. BCAR: Bilinear Cross-Attention with Feature Replacement**

**2.1. Architecture and Design**

BCAR extends the Bilateral Cross-Attention (BCAA) framework but imposes a replacement constraint that removes all independent chemical features from the final representation. As in BCAA, linker and payload embeddings are fused through a bidirectional cross-attention mechanism. The key modification is that the original independent embeddings (CLS-level linker and payload features) are discarded rather than concatenated with the interaction features. Consequently, the model forwards only the cross-attended interaction representation into the terminal MLP, such that the final feature vector equals the cross-attention output. This design slightly reduces the parameter count relative to BCAA (5.2M vs. 5.3M parameters) because the MLP receives a lower-dimensional input, though it remains larger than ABFormer (4.7M). Conceptually, BCAR operationalizes an “interaction-dominance” hypothesis, positing that linker–payload interactions alone determine efficacy and that independent molecular features are unnecessary or even detrimental.

**2.2. Training Dynamics**

Under Seed 1, BCAR exhibits a deceptively strong training profile on the random split. It attains a peak validation AUC of 0.862 (ABFormer: 0.842) and a minimum validation loss of 0.290 (ABFormer: 0.277). This performance, combined with the later collapse under LOPO evaluation, indicates that the model memorizes interaction signatures specific to the random split rather than acquiring transferable rules. The behaviour is consistent with overfitting driven by an overreliance on the interaction module.

**2.3. LP Performance (15 Seeds)**

Across 15 seeds, BCAR shows degraded generalization relative to ABFormer. Its mean MCC is 0.630 ± 0.15 (ABFormer: 0.696 ± 0.09), specificity declines markedly from 0.914 ± 0.12 to 0.820 ± 0.21, and accuracy drops from 0.844 ± 0.03 to 0.819 ± 0.06. AUC remains broadly comparable (0.888 ± 0.01 vs. 0.898 ± 0.01). The pronounced loss in specificity and the large variance indicate systematic instability: without access to independent molecular representations, the model frequently misclassifies negatives as positives, reflecting a reliance on an interaction module that is difficult to learn robustly in a small-data setting.

**2.4. Benchmark Analysis**

The failure on negative controls illustrates the miscalibration. For Sample 3 (true negative), BCAR outputs 0.519, resulting in a false positive, whereas ABFormer correctly assigns 0.479. For Sample 22 (true negative), BCAR predicts 0.496 (borderline), in contrast to ABFormer’s clearly negative 0.449. These behaviours demonstrate that BCAR systematically shifts non-binding compounds toward the decision boundary or beyond it, producing an overly aggressive “Active” bias.

**2.5. Root Cause and Conclusion**

The underlying failure mechanism is feature starvation. In ABFormer, independent linker and payload encoders supply a stable baseline characterizing intrinsic molecular properties, while cross-attention modulates these properties through interaction signals. BCAR eliminates this baseline and forces the model to reconstruct molecular identity exclusively from a high-complexity interaction matrix. Given the limited dataset (~340 samples), the interaction-only representation is insufficiently informative, leading to unstable decision boundaries and erroneous inferences of compatibility.

Overall, the findings underscore that redundancy enhances robustness: in low-data regimes, independent molecular features should complement, not be replaced by, complex interaction modules.

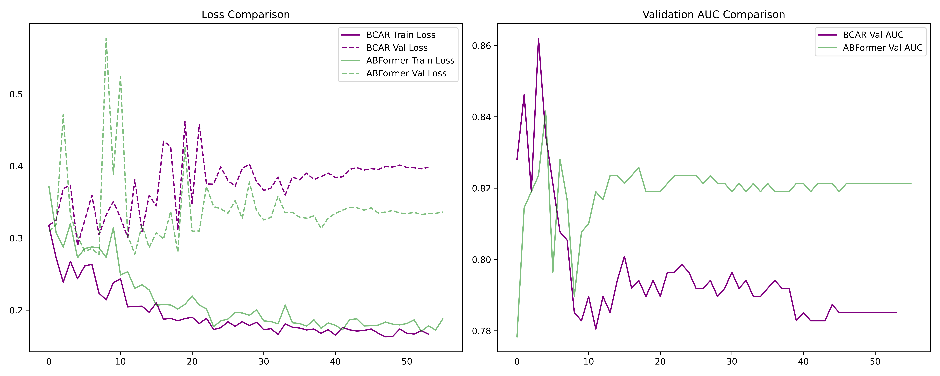

**Supplementary Figure 2.** Training and validation performance comparison between BCAR and ABFormer. The left panel illustrates train and validation loss trajectories, where solid and dashed purple curves represent BCAR training and validation loss, respectively, and solid and dashed green curves represent ABFormer training and validation loss. The right panel shows validation AUC evolution across epochs, with BCAR plotted in purple and ABFormer in green.

**3. BLICAF: Bilinear Gated Fusion with LLM Encoders**

**3.1. Architecture and Design**

BLICAF introduces a full architectural replacement, modifying both the molecular encoders and the fusion strategy simultaneously. In contrast to baselines that vary only a single component, BLICAF substitutes the specialized FG-BERT encoders with large chemical language models and replaces simple concatenation with a bilinear gated interaction network.

For the linker, the model employs MegaMolBART, a BART-derived transformer pre-trained on ZINC, while the payload is encoded with MolFormer, a linear-attention transformer trained on large-scale PubChem and ZINC corpora. The interaction module implements a Bilinear Gated Fusion network with three pathways capturing linker–payload, linker–antibody, and payload–conjugate relationships. The core operator is a bilinear gated unit that computes a multiplicative interaction modulated by a sigmoid gate, and this unit is applied recursively to generate the final drug-level embedding. The total parameter count reaches 5,618,754—approximately 20% higher than ABFormer—owing primarily to the dense projection matrices required to align the high-dimensional LLM embeddings to the bilinear fusion space.

Mathematical Formulation:The core operator is the Bilinear Gated unit $B(a,b)$, which learns a multiplicative interaction between features, regulated by a sigmoid gate $\sigma$:

$B(a,b)=\sigma(W_{g}(a\odot b))\odot W_{o}(a\odot b)$ , (1)

This operator is applied recursively to build the final drug embedding:

$h_{LP}=[B(h_{Li},h_{P}),B(h_{Li},h_{P})]$ , (2)

$$h_{Drug}=f_{Drug}\left( \left[ h_{LP},h_{liCtx},h_{PCtx},h_{d} \right] \right) , (3)$$

**3.2. Training Dynamics**

Under Seed 1, the training trajectory is dominated by the “LLM effect”: rapid convergence appears immediately due to the rich, linearly separable molecular representations produced by the pre-trained encoders. Although BLICAF attains a peak validation AUC of 0.830, comparable to ABFormer’s 0.842, it fails to achieve a correspondingly low validation loss. The behaviour indicates that the model does not acquire new chemical structure–activity patterns from the ADC dataset but instead exploits pre-existing clustering inherent to the MolFormer and MegaMolBART embedding spaces, with the bilinear gates amplifying these latent groupings.

**3.3. LP Performance (15 Seeds)**

Across LP evaluation, BLICAF yields the highest mean AUC of all models (0.936 ± 0.02 compared with ABFormer’s 0.898 ± 0.01) and slightly higher accuracy (0.855 ± 0.04 versus 0.844 ± 0.03). However, this apparent superiority masks a detrimental trade-off: sensitivity increases (0.864 ± 0.07 versus 0.802 ± 0.02) at the expense of specificity, which declines substantially from 0.914 ± 0.12 to 0.839 ± 0.18. The model thus excels at identifying positives while failing to reject structurally plausible but inactive candidates, indicating that the increased AUC arises from aggressive positive recall rather than balanced discrimination.

**3.4. Benchmark Analysis**

The specificity collapse is evident in the negative-control predictions. For Sample 3 (true negative), ABFormer outputs 0.479, whereas BLICAF assigns 0.968, a clear false positive; for Sample 22 (true negative), ABFormer predicts 0.449 while BLICAF again outputs 0.959. Such extreme misclassifications reflect a severe domain-shift failure: BLICAF produces near-saturated activity probabilities across samples and behaves more like a detector of potent payload chemistry than an ADC efficacy model.

**3.5. Root Cause and Conclusion**

The principal failure mechanism is a “super-detector” effect arising from the combination of highly expressive chemical LLM encoders and the amplifying bilinear gate. MegaMolBART and MolFormer encode extensive prior knowledge about toxicity, drug-likeness, and other intrinsic molecular properties learned from millions of molecules. The bilinear operator magnifies these signals, enabling a shortcut in which the model predicts activity whenever the payload resembles a toxin and the linker appears cleavable, largely ignoring the antibody-dependent biological context fundamental to ADC efficacy.

In contrast, ABFormer’s FG-BERT encoders capture functional groups without embedding broad toxicity priors, forcing the model to incorporate antibody–antigen context in its predictions. Consequently, BLICAF—despite its chemical sophistication—overfits to payload potency and misjudges systemic performance.

Overall, these results demonstrate that in context-dependent biological prediction tasks, excessively powerful chemical encoders can become distractors, leading models to prioritize intrinsic molecular properties over the biologically critical conjugation and delivery mechanisms.

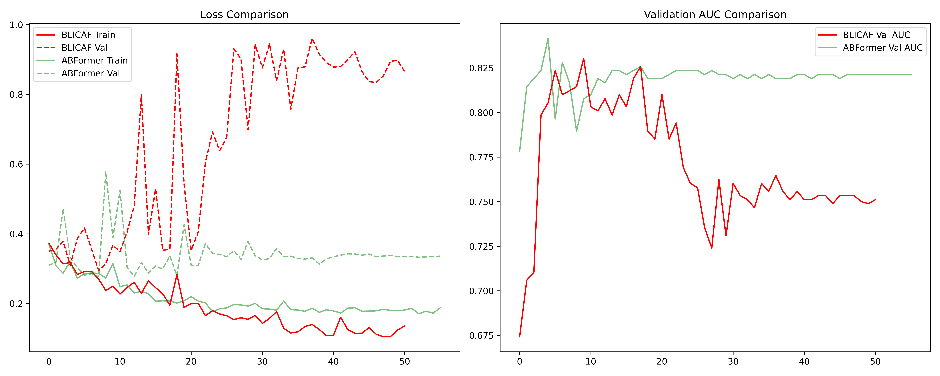

**Supplementary Figure 3.** Comparison of training dynamics between BLICAF and ABFormer. The left panel shows training and validation loss curves, with solid red and dashed red lines representing BLICAF training and validation loss, and solid green and dashed green lines representing ABFormer training and validation loss. The right panel illustrates validation AUC trajectories, where BLICAF is plotted in red and ABFormer in green, highlighting the performance divergence across epochs.

**4. MLP: LLM Encoders with Simple Concatenation**

**4.1. Architecture and Design**

This model replaces the compact, domain-specific FG-BERT encoders with large chemical language models and simultaneously simplifies the fusion mechanism to a raw concatenation strategy. Payloads are encoded using MolFormer, which produces 768-dimensional representations through a linear-attention transformer trained on PubChem and ZINC, while linkers are encoded using MegaMolBART, a BART-based model pre-trained on ZINC that yields 512-dimensional embeddings. These LLM-derived features are concatenated directly with antibody, antigen, light-chain, and DAR inputs, forming an exceptionally high-dimensional vector that feeds into a deep multilayer perceptron classifier. The resulting architecture contains approximately 7.2 million parameters—an increase of about 2.5 million over ABFormer—due to the need for large fully connected layers capable of processing the inflated input space. The underlying hypothesis is that state-of-the-art molecular LLMs, trained on millions of compounds, might provide sufficiently rich representations to improve ADC activity prediction without requiring complex interaction modeling.

**4.2. Training Dynamics**

The Seed 1 training trajectory demonstrates a clear convergence failure. Whereas ABFormer reaches a minimum validation loss of 0.27, the MLP stabilizes at a much higher value of 1.52. Since random guessing typically yields a validation loss near 0.69, this result reflects not mere underfitting but confident misclassification: the model memorizes training patterns intensely yet generalizes poorly, producing high-certainty errors on unseen samples. The best validation AUC of only 0.7919 reinforces this interpretation and highlights the absence of robust structure–activity learning.

**4.3. LP Performance (15 Seeds)**

Across LP evaluation, the MLP performs worst among all tested architectures. The mean MCC falls to 0.485 ± 0.11 (ABFormer: 0.696 ± 0.09), specificity drops sharply to 0.682 ± 0.19 (ABFormer: 0.914 ± 0.12), and accuracy declines to 0.752 ± 0.05. Even the AUC, at 0.868 ± 0.03, remains below ABFormer’s 0.898 ± 0.01. The pronounced specificity deficit indicates that the model predicts the “Active” class by default, failing to capture the subtle signals required to differentiate non-binders from true positives. This behaviour is consistent with high-dimensional noise obscuring informative negative cues.

**4.4. Benchmark Analysis**

The model’s collapse becomes evident when evaluated on negative controls. For Sample 3, a true negative, ABFormer assigns a probability of 0.479, whereas the MLP outputs 0.939; for Sample 22, ABFormer predicts 0.449 while the MLP yields 0.949. These errors occur with high confidence and illustrate severe miscalibration. Moreover, prediction variance across seeds is extreme—some samples oscillate between near-zero and near-one values—revealing that the model behaves as a high-variance estimator biased toward positive outputs rather than a stable classifier suitable for external application.

**4.5. Root Cause and Conclusion**

The underlying cause is a combination of dimensionality explosion and parameter overgrowth. MolFormer and MegaMolBART encode broad chemical priors learned from millions of molecules, but when transferred to a dataset of only ~340 ADC instances, their high-dimensional embeddings inject substantial variance without providing task-specific structure. The 7.2-million-parameter MLP amplifies this issue: the network has sufficient capacity to memorize the training set but lacks the regularization and data volume needed to generalize. The biological context encoded in the AntiBinder features is effectively drowned out by the large chemical vectors, leading the model to ignore the antibody-dependent determinants of ADC efficacy.

Taken together, these results reaffirm that model capacity must be aligned with dataset scale. For small, mechanistically specialized datasets, compact encoders like FG-BERT outperform massive, general-purpose LLMs; larger models are not merely unnecessary but actively harmful, introducing noise that overwhelms the signal.

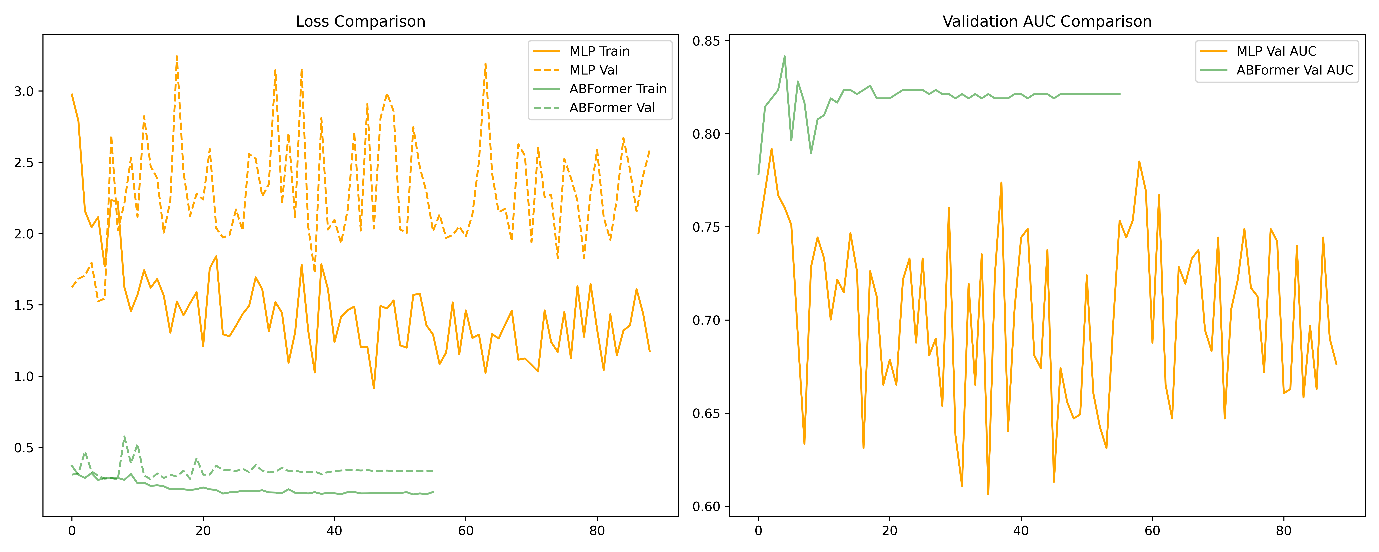

**Supplementary Figure 4.** Training and validation performance comparison between a baseline MLP model and ABFormer. The left panel presents training and validation loss curves, with solid and dashed orange lines indicating MLP training and validation loss, and solid and dashed green lines indicating ABFormer training and validation loss. The right panel shows validation AUC trajectories, where the MLP model is plotted in orange and ABFormer in green, demonstrating the stability and improved predictive consistency of ABFormer across epochs.

**5. MLP_MMB: Dual MegaMolBART Encoders with Simple Concatenation**

**5.1. Architecture and Design**

The MLP_MMB model replaces the domain-specific FG-BERT encoders with a single, high-capacity chemical language model for both small-molecule components and simultaneously simplifies the fusion strategy to raw feature concatenation. Specifically, both the linker and payload are encoded using MegaMolBART, each producing 512-dimensional representations derived from a BART-based transformer pre-trained on large chemical corpora. These embeddings are concatenated with antibody, antigen, and DAR features and passed directly to a deep multilayer perceptron classifier. The resulting architecture contains approximately 6.98 million parameters, compared with 4.7 million for ABFormer, with the increase driven primarily by the large input projection layer required to accommodate the concatenated LLM-derived features. The motivating hypothesis was that using the same high-capacity encoder for both chemical components, thereby enforcing a shared latent chemical space, might improve performance relative to mixed or simpler encoders.

**5.2. Training Dynamics**

Training behaviour under Seed 1 indicates complete convergence failure. While ABFormer reaches a minimum validation loss of 0.27, MLP_MMB attains a minimum validation loss of 2.43. For a binary classification task, a validation loss exceeding 2.0 is indicative of a fundamentally unstable model. This suggests that the optimizer is unable to navigate the loss landscape induced by the extremely high-dimensional input space, likely producing poorly calibrated or oscillatory logits that incur large penalties during validation.

**5.3. LP Performance (15 Seeds)**

LP evaluation confirms that MLP_MMB performs near the level of random guessing. The mean MCC is 0.484 ± 0.19, far below ABFormer’s 0.696 ± 0.09, and specificity collapses to 0.678 ± 0.21 compared with 0.914 ± 0.12. Accuracy is reduced to 0.757 ± 0.08, and AUC declines to 0.844 ± 0.08. The large standard deviations across all metrics reveal severe instability: some random seeds occasionally converge to weak local minima, while others diverge almost entirely, underscoring the absence of a reliable decision boundary.

**5.4. Benchmark Analysis**

The failure mode is further exposed by negative-control predictions. For Sample 3, a true negative, ABFormer outputs 0.479, whereas MLP_MMB assigns a probability of 0.902. Similarly, for Sample 22, ABFormer predicts 0.449 while MLP_MMB outputs 0.939. Across benchmarks, the model consistently produces probabilities in the 0.80–0.95 range, indicating a collapse toward predicting the majority “Active” class irrespective of input features. This behaviour reflects a loss of calibration and effective disregard for discriminative signals.

**5.5. Root Cause and Conclusion**

The dominant failure mechanism is latent space incompatibility coupled with dimensional overload. MegaMolBART embeddings encode chemically rich but biologically agnostic information. When two such high-variance chemical vectors are concatenated with biological features, the combined representation becomes dominated by chemical variance, overwhelming the antibody-dependent context critical for ADC efficacy. With only ~340 training samples, the optimizer cannot identify a separating hyperplane in a space exceeding 6,000 dimensions and instead converges to the trivial solution of predicting the dataset prior.
These results reinforce that domain specificity is essential: deploying a general-purpose chemical LLM for both linker and payload not only fails to improve performance but introduces sufficient noise to destabilize training entirely.

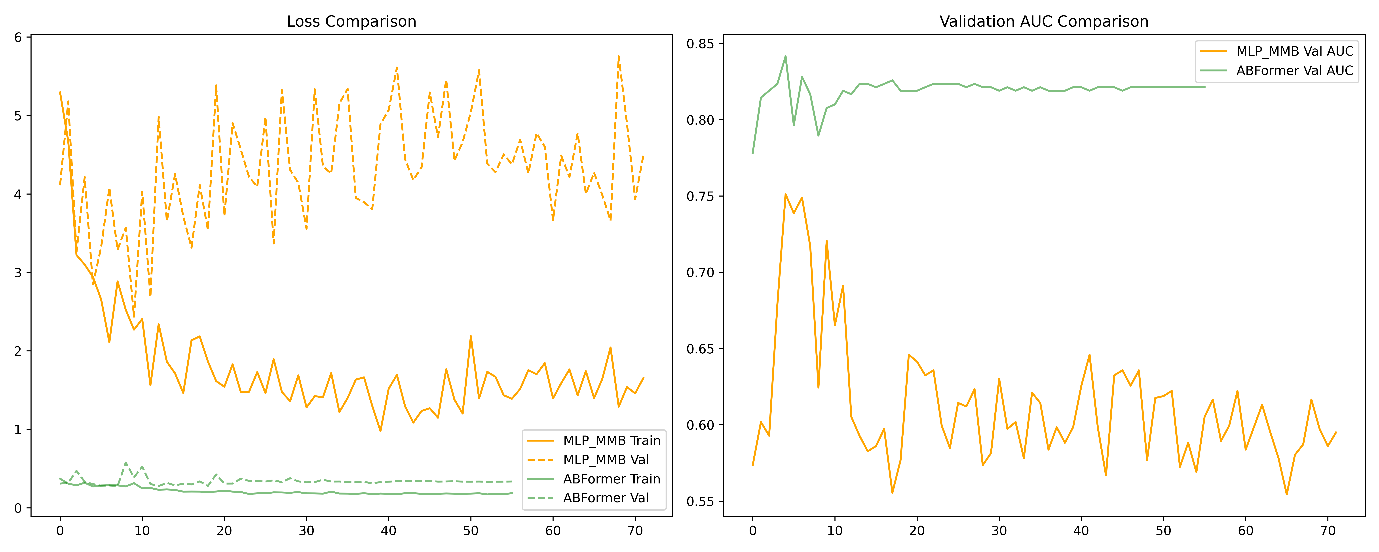

**Supplementary Figure 5.** Comparison of learning dynamics between the MLP_MMB baseline and ABFormer. The left panel depicts training and validation loss curves, with solid and dashed orange lines corresponding to MLP_MMB training and validation loss, and solid and dashed green lines corresponding to ABFormer training and validation loss. The right panel shows validation AUC trajectories, where MLP_MMB is plotted in orange and ABFormer in green, demonstrating the substantially higher stability and predictive performance achieved by ABFormer across epochs.

**6. MMF: Attention-Based Multimodal Fusion**

**6.1. Architecture and Design**

MMF replaces standard molecular encoders with large chemical language models and employs a unified self-attention–based fusion mechanism. Linkers are represented using MegaMolBART and payloads with MolFormer, while antibody heavy-chain, light-chain and antigen features form additional modalities. All five components are projected into a shared latent space and treated as tokens in a short multimodal sequence. A single self-attention layer assigns modality-specific fusion weights, producing a weighted sum that serves as the final representation. This strategy yields a relatively lightweight architecture, adding roughly 2.5 million parameters over ABFormer by compressing all high-dimensional embeddings to 256 dimensions before fusion. The design aims to determine automatically whether the antibody–antigen interface (specificity) or the payload chemistry (potency) should dominate prediction, without embedding explicit mechanistic constraints.

**6.2. Training Dynamics**

Under Seed 1, MMF displays rapid and aggressive convergence. It achieves a validation AUC of 0.867, exceeding ABFormer’s 0.842, and reduces training loss efficiently. The pattern suggests that the attention layer rapidly identifies correlations in the training set—most notably that MolFormer’s toxin-like embeddings align strongly with positive labels—leading the model to allocate disproportionate attention to the chemical modality.

**6.3. LP Performance (15 Seeds)**

In LP evaluation, MMF attains the highest mean AUC among all models (0.923 ± 0.03), but this apparent superiority masks weakened classification behaviour. Specificity drops to 0.878 ± 0.12 from ABFormer’s 0.914 ± 0.12, whereas sensitivity rises modestly (0.818 ± 0.09 versus 0.802 ± 0.02). This metric profile—elevated AUC but reduced specificity—is indicative of a model that systematically shifts its probability scale upward. Active compounds are pushed toward 0.99, while inactive compounds shift upward into the 0.50–0.70 range, preserving ranking but degrading threshold-based discrimination.

**6.4. Benchmark Analysis**

The benchmark results reveal the underlying fragility of unconstrained self-attention. For Sample 3 (a true negative), MMF assigns 0.981, whereas ABFormer correctly predicts 0.479. For Sample 22, MMF outputs 0.897 compared with ABFormer’s 0.449. Moreover, MMF fails on Sample 21, a true positive, assigning only 0.350. Across the benchmark, the model produces scores exceeding 0.94 for most compounds, including negative controls, indicating severe miscalibration. Notably, the model assigns the negative control Sample 3 a higher probability than multiple true positives, demonstrating that payload-specific chemical features dominate the attention distribution.

**6.5. Root Cause and Conclusion**

The failure originates from an imbalance in representational “loudness” between chemical and biological features. MolFormer encodings of cytotoxic payloads possess high-magnitude, tightly clustered patterns, whereas antibody and antigen embeddings capture subtle structural compatibilities that are lower in amplitude. In a self-attention architecture, dominant features systematically attract greater weights, enabling the chemical modality to eclipse the biological modalities. Consequently, MMF learns to predict activity primarily from payload potency, disregarding the antibody-dependent delivery mechanism essential to ADC efficacy.
These observations underscore the limitations of relying solely on soft attention in small biological datasets. Without architectural constraints enforcing biologically meaningful interactions, the model gravitates toward the easiest signal—payload toxicity—rather than the causal determinants of ADC activity. ABFormer succeeds precisely because its AntiBinder stream structurally enforces antibody–antigen reasoning and prevents the molecular signal from bypassing that pathway.

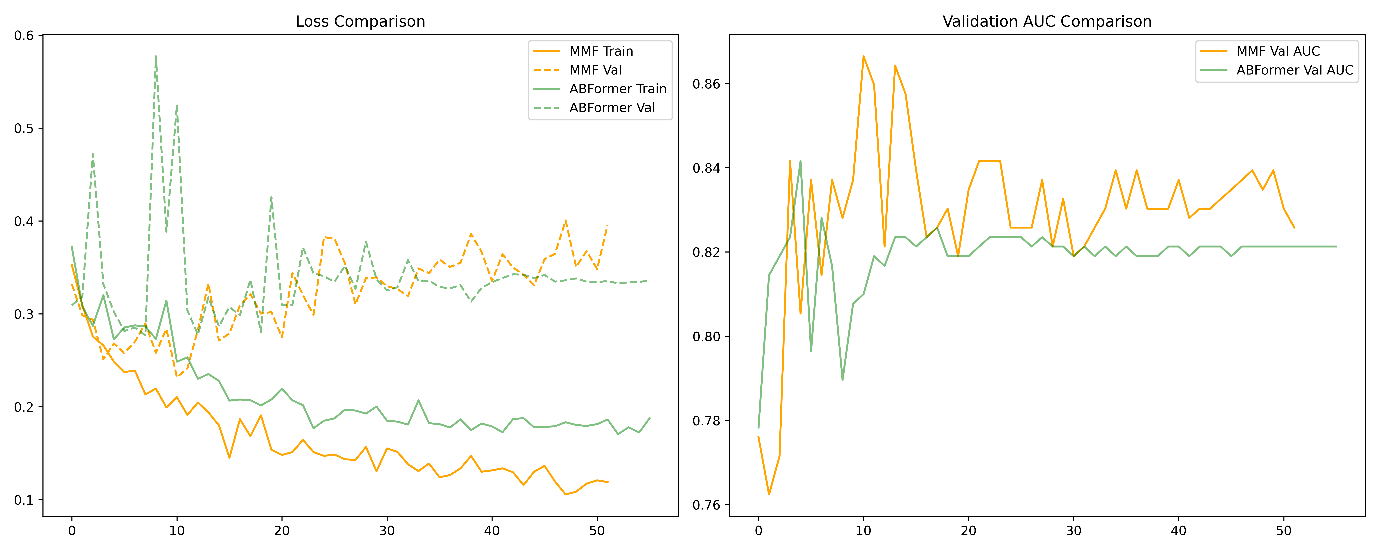

**Supplementary Figure 6.** Training and validation performance comparison between the MMF model and ABFormer. The left panel presents train and validation loss trajectories, where solid and dashed orange lines denote MMF training and validation loss, and solid and dashed green lines denote ABFormer training and validation loss. The right panel shows validation AUC evolution across epochs, with MMF plotted in orange and ABFormer in green, highlighting the comparative stability and predictive behaviour of both models.

**Supplementary Table T1:** Comparative Performance on Random Split (LP), reported as mean ± standard deviation over 15 seeds for baseline ML models, ADCNet, and ABFormer.

| **Model** | **SE** | **SP** | **MCC** | **ACC** | **AUC** | **F1** | **BA** | **PRAUC** | **PPV** | **NPV** |
| --- | --- | --- | --- | --- | --- | --- | --- | --- | --- | --- |
| **ADCNet** | 0.8664±0.0802 | 0.6219±0.1567 | 0.5115±0.1376 | 0.7788±0.0630 | 0.8457±0.0627 | 0.8301±0.0558 | 0.7442±0.0764 | 0.8907±0.0570 | 0.8040±0.0757 | 0.7375±0.1223 |
| **ABFormer** | 0.6717±0.1191 | **0.8574±0.0858** | 0.5122±0.1072 | 0.7333±0.0741 | 0.8686±0.0489 | 0.7623±0.0672 | 0.7646±0.0457 | 0.9268±0.0277 | **0.9018±0.0497** | 0.5951±0.1368 |
| **XGB (MACCS)** | 0.9103±0.0218 | 0.7511±0.0890 | 0.6758±0.0675 | 0.8561±0.0281 | 0.9068±0.0201 | 0.8932±0.0187 | 0.8307±0.0415 | 0.9496±0.0139 | 0.8779±0.0365 | 0.8133±0.0340 |
| **SVM (MACCS)** | 0.8943±0.0275 | 0.7867±0.0276 | 0.6837±0.0338 | 0.8576±0.0160 | 0.8352±0.0190 | 0.8920±0.0133 | **0.8405±0.0141** | 0.8618±0.0316 | 0.8904±0.0112 | 0.7963±0.0440 |
| **RF (MACCS)** | **0.9333±0.0205** | 0.6622±0.0469 | 0.6365±0.0581 | 0.8409±0.0243 | 0.8962±0.0129 | 0.8855±0.0173 | 0.7978±0.0287 | 0.9450±0.0089 | 0.8426±0.0199 | **0.8379±0.0476** |
| **LR (MACCS)** | 0.8069±0.0408 | 0.7733±0.0607 | 0.5660±0.0312 | 0.7955±0.0149 | 0.9238±0.0066 | 0.8383±0.0150 | 0.7901±0.0174 | 0.9613±0.0040 | 0.8748±0.0269 | 0.6775±0.0347 |
| **XGB (Morgan)** | 0.9241±0.0350 | 0.7467±0.0843 | **0.6921±0.0926** | **0.8636±0.0403** | **0.9298±0.0116** | **0.8994±0.0292** | 0.8354±0.0488 | **0.9650±0.0060** | 0.8768±0.0362 | 0.6775±0.0347 |
| **SVM (Morgan)** | 0.8575±0.0503 | 0.7733±0.0338 | 0.6259±0.0813 | 0.8288±0.0411 | 0.8607±0.0216 | 0.8679±0.0347 | 0.8154±0.0378 | 0.9019±0.0210 | 0.8793±0.0206 | 0.7418±0.0692 |
| **RF (Morgan)** | 0.9126±0.0178 | 0.7378±0.0733 | 0.6679±0.0503 | 0.8530±0.0208 | 0.9163±0.0164 | 0.8913±0.0135 | 0.8252±0.0325 | 0.9582±0.0092 | 0.8718±0.0293 | 0.8147±0.0252 |
| **LR (Morgan)** | 0.7747±0.0535 | 0.8000±0.0000 | 0.5536±0.0571 | 0.7833±0.0353 | 0.9195±0.0065 | 0.8241±0.0340 | 0.7874±0.0268 | 0.9542±0.0048 | 0.8818±0.0074 | 0.6516±0.0531 |

The best performing model for each metric is highlighted in bold.

**Supplementary Table T2:** Comparative Performance on Leave-Pair-Out (LP) of Ablation Variants and ABFormer, reported as mean ± standard deviation over 15 seeds.

| **Model** | **test_se** | **test_sp** | **test_mcc** | **test_acc** | **test_auc** | **test_f1** | **test_ba** | **test_prauc** | **test_ppv** | **test_npv** |
| --- | --- | --- | --- | --- | --- | --- | --- | --- | --- | --- |
| **ABFormer** | 0.8023 ± 0.0243 | 0.9137 ± 0.1194 | 0.6964 ± 0.0895 | 0.8435 ± 0.0331 | 0.8983 ± 0.0061 | 0.8668 ± 0.0218 | 0.8580 ± 0.0505 | 0.9520 ± 0.0024 | 0.9475 ± 0.0700 | 0.7299 ± 0.0167 |
| **w/o AACMACCS** | 0.8138 ± 0.0387 | 0.8588 ± 0.1871 | 0.6612 ± 0.1288 | 0.8304 ± 0.0474 | 0.8902 ± 0.0055 | 0.8600 ± 0.0275 | 0.8363 ± 0.0759 | 0.9496 ± 0.0027 | 0.9222 ± 0.0965 | 0.7287 ± 0.0170 |
| **w/o AAC** | 0.8092 ± 0.0388 | 0.8079 ± 0.1165 | 0.6080 ± 0.0708 | 0.8087 ± 0.0221 | 0.8972 ± 0.0040 | 0.8425 ± 0.0119 | 0.8085 ± 0.0408 | 0.9514 ± 0.0018 | 0.8849 ± 0.0671 | 0.7144 ± 0.0179 |
| **w/o DAR** | 0.8115 ± 0.0430 | 0.8588 ± 0.2118 | 0.6597 ± 0.1440 | 0.8290 ± 0.0532 | 0.8956 ± 0.0052 | 0.8591 ± 0.0299 | 0.8352 ± 0.0857 | 0.9506 ± 0.0022 | 0.9253 ± 0.1042 | 0.7256 ± 0.0169 |
| **w/o MACCS** | 0.8069 ± 0.0565 | 0.8510 ± 0.2424 | 0.6505 ± 0.1589 | 0.8232 ± 0.0580 | 0.8928 ± 0.0054 | 0.8546 ± 0.0316 | 0.8290 ± 0.0955 | 0.9510 ± 0.0026 | 0.9259 ± 0.1163 | 0.7197 ± 0.0213 |
| **w/o Chem. Embeds** | 0.8460 ± 0.0449 | 0.7647 ± 0.1792 | 0.6142 ± 0.1131 | 0.8159 ± 0.0410 | 0.8975 ± 0.0035 | 0.8543 ± 0.0231 | 0.8054 ± 0.0691 | 0.9513 ± 0.0016 | 0.8725 ± 0.0920 | 0.7464 ± 0.0237 |
| **w/o Antigen** | 0.8184 ± 0.0545 | 0.8353 ± 0.2052 | 0.6485 ± 0.1250 | 0.8246 ± 0.0438 | 0.8967 ± 0.0044 | 0.8565 ± 0.0231 | 0.8269 ± 0.0768 | 0.9512 ± 0.0020 | 0.9128 ± 0.1044 | 0.7321 ± 0.0177 |
| **Zeroed Antibinder Embeddings** | 0.5058 ± 0.0998 | 0.7177 ± 0.0456 | 0.2199 ± 0.0738 | 0.5841 ± 0.0538 | 0.7164 ± 0.0097 | 0.6000 ± 0.0803 | 0.6117 ± 0.0395 | 0.7475 ± 0.0247 | 0.7523 ± 0.0290 | 0.4645 ± 0.0459 |

**Supplementary Table T3**: Comparative Performance on Leave-Pair-Out (LP) of Alternative Architectures, reported as mean ± standard deviation over 15 seeds.

| **Model** | **test_se** | **test_sp** | **test_mcc** | **test_acc** | **test_auc** | **test_f1** | **test_ba** | **test_prauc** | **test_ppv** | **test_npv** |
| --- | --- | --- | --- | --- | --- | --- | --- | --- | --- | --- |
| **MMF** | 0.8184 ± 0.0859 | 0.8784 ± 0.1227 | 0.6886 ± 0.0692 | 0.8406 ± 0.0364 | 0.9229 ± 0.0334 | 0.8647 ± 0.0375 | 0.8484 ± 0.0410 | 0.9593 ± 0.0213 | 0.9292 ± 0.0663 | 0.7521 ± 0.0746 |
| **BCAA** | 0.8092 ± 0.0595 | 0.8196 ± 0.1449 | 0.6213 ± 0.0945 | 0.8130 ± 0.0356 | 0.8867 ± 0.0064 | 0.8453 ± 0.0264 | 0.8144 ± 0.0539 | 0.9478 ± 0.0033 | 0.8949 ± 0.0792 | 0.7192 ± 0.0417 |
| **MMFmega** | 0.7954 ± 0.0819 | **0.9412 ± 0.0832** | **0.7182 ± 0.0766** | 0.8493 ± 0.0461 | 0.9345 ± 0.0220 | 0.8675 ± 0.0479 | **0.8683 ± 0.0426** | 0.9662 ± 0.0145 | **0.9631 ± 0.0485** | 0.7372 ± 0.0660 |
| **BCAR** | 0.8184 ± 0.0401 | 0.8196 ± 0.2126 | 0.6301 ± 0.1517 | 0.8188 ± 0.0579 | 0.8880 ± 0.0094 | 0.8530 ± 0.0341 | 0.8190 ± 0.0894 | 0.9479 ± 0.0047 | 0.9019 ± 0.1022 | 0.7222 ± 0.0297 |
| **MLPmmb** | 0.8023 ± 0.0743 | 0.6784 ± 0.2096 | 0.4841 ± 0.1900 | 0.7565 ± 0.0810 | 0.8435 ± 0.0812 | 0.8072 ± 0.0614 | 0.7404 ± 0.1030 | 0.9121 ± 0.0558 | 0.8220 ± 0.1054 | 0.6670 ± 0.0913 |
| **BLICAF** | **0.8644 ± 0.0707** | 0.8392 ± 0.1826 | 0.7070 ± 0.1018 | **0.8551 ± 0.0416** | **0.9359 ± 0.0170** | **0.8833 ± 0.0281** | 0.8518 ± 0.0665 | **0.9689 ± 0.0094** | 0.9156 ± 0.0881 | **0.7976 ± 0.0725** |

The best performing model for each metric is highlighted in bold.

**Supplementary Table T4:** Mean predicted probability ± standard deviation on the independent benchmark using 15-seed inference of Alternate Architectures. Bold values indicate negative samples.

| **Sample** | **True Label** | **BCAR** | **BCAA** | **BLI CAF** | **MLP** | **MLP_mmb** | **MMF** |
| --- | --- | --- | --- | --- | --- | --- | --- |
| **1** | 1 | 0.8917 ± 0.0263 | 0.8837 ± 0.0353 | 0.5384 ± 0.3888 | 0.5710 ± 0.4357 | 0.8713 ± 0.3114 | 0.9402 ± 0.0357 |
| **2** | 1 | 0.7632 ± 0.0538 | 0.7351 ± 0.0740 | 0.9566 ± 0.0267 | 0.9859 ± 0.0401 | 0.9953 ± 0.0134 | 0.9585 ± 0.0326 |
| **3** | **0** | **0.5194 ± 0.0766** | **0.4739 ± 0.0782** | **0.9678 ± 0.0109** | **0.9393 ± 0.2255** | **0.9017 ± 0.2541** | **0.9814 ± 0.0125** |
| **4** | 1 | 0.6548 ± 0.0637 | 0.6234 ± 0.0717 | 0.9080 ± 0.0987 | 0.3872 ± 0.4497 | 0.3155 ± 0.4363 | 0.5005 ± 0.3060 |
| **5** | 1 | 0.6759 ± 0.0592 | 0.6379 ± 0.0742 | 0.9659 ± 0.0152 | 0.3287 ± 0.4109 | 0.3163 ± 0.4309 | 0.5997 ± 0.2323 |
| **6** | 1 | 0.6035 ± 0.0494 | 0.6073 ± 0.0743 | 0.9563 ± 0.0401 | 0.3252 ± 0.4042 | 0.2931 ± 0.4087 | 0.6085 ± 0.2982 |
| **7** | 1 | 0.8998 ± 0.0384 | 0.8778 ± 0.0381 | 0.9691 ± 0.0151 | 0.9666 ± 0.1112 | 0.8065 ± 0.3752 | 0.7644 ± 0.2076 |
| **8** | 1 | 0.9065 ± 0.0360 | 0.8795 ± 0.0381 | 0.9649 ± 0.0195 | 0.9497 ± 0.1834 | 0.8045 ± 0.3791 | 0.6223 ± 0.2662 |
| **9** | 1 | 0.9114 ± 0.0252 | 0.8803 ± 0.0372 | 0.9641 ± 0.0262 | 0.9507 ± 0.1793 | 0.8048 ± 0.3785 | 0.5342 ± 0.2625 |
| **10** | 1 | 0.9080 ± 0.0270 | 0.8798 ± 0.0377 | 0.9691 ± 0.0125 | 0.9529 ± 0.1687 | 0.8048 ± 0.3786 | 0.7956 ± 0.1873 |
| **11** | 1 | 0.9150 ± 0.0232 | 0.8812 ± 0.0367 | 0.9682 ± 0.0135 | 0.9522 ± 0.1678 | 0.8065 ± 0.3753 | 0.7331 ± 0.2289 |
| **12** | 1 | 0.9037 ± 0.0315 | 0.8793 ± 0.0383 | 0.9628 ± 0.0255 | 0.9446 ± 0.1932 | 0.8043 ± 0.3796 | 0.6174 ± 0.2329 |
| **13** | 1 | 0.9013 ± 0.0343 | 0.8788 ± 0.0383 | 0.9607 ± 0.0287 | 0.9413 ± 0.2057 | 0.8043 ± 0.3795 | 0.5463 ± 0.2458 |
| **14** | 1 | 0.9081 ± 0.0275 | 0.8799 ± 0.0374 | 0.9589 ± 0.0306 | 0.9385 ± 0.2058 | 0.8047 ± 0.3788 | 0.6588 ± 0.2281 |
| **15** | 1 | 0.8885 ± 0.0364 | 0.8758 ± 0.0382 | 0.9701 ± 0.0112 | 0.9752 ± 0.0837 | 0.8063 ± 0.3758 | 0.7744 ± 0.1983 |
| **16** | 1 | 0.8888 ± 0.0361 | 0.8756 ± 0.0381 | 0.9716 ± 0.0076 | 0.9798 ± 0.0682 | 0.8067 ± 0.3749 | 0.8465 ± 0.1348 |
| **17** | 1 | 0.9048 ± 0.0289 | 0.8801 ± 0.0373 | 0.9623 ± 0.0232 | 0.9427 ± 0.2061 | 0.8044 ± 0.3794 | 0.6575 ± 0.2183 |
| **18** | 1 | 0.8842 ± 0.0326 | 0.8724 ± 0.0377 | 0.9715 ± 0.0076 | 0.9780 ± 0.0800 | 0.8078 ± 0.3729 | 0.8293 ± 0.1979 |
| **19** | 1 | 0.8874 ± 0.0321 | 0.8730 ± 0.0380 | 0.9682 ± 0.0147 | 0.9823 ± 0.0577 | 0.8070 ± 0.3744 | 0.8504 ± 0.1394 |
| **20** | 1 | 0.9474 ± 0.0194 | 0.9307 ± 0.0246 | 0.8258 ± 0.2039 | 0.8873 ± 0.3150 | 0.8485 ± 0.3421 | 0.7824 ± 0.1916 |
| **21** | 1 | 0.9928 ± 0.0039 | 0.9874 ± 0.0057 | 0.9472 ± 0.0461 | 0.9517 ± 0.1788 | 0.9569 ± 0.1219 | 0.3503 ± 0.2492 |
| **22** | **0** | **0.4959 ± 0.0761** | **0.4844 ± 0.0822** | **0.959 ± 0.076** | **0.9490 ± 0843** | **0.9387 ± 0.0549** | **0.8967 ± 0.0968** |
